# Reducing Manual Labour in Forensic Microtrace Recognition with Deep Learning

**DOI:** 10.1101/2025.05.09.653056

**Authors:** Gerben Rijpkema, Dylan Kalisvaart, Serafim Korovin, Daniel Spengler, Anna Pals, Jaap van der Weerd, Carlas Smith

**Affiliations:** Delft Center for Systems and Control, Delft University of Technology, Delft, The Netherlands; Netherlands Forensic Institute (NFI), The Hague, The Netherlands

## Abstract

Forensic microtrace investigation relies on time- and labor-intensive microscopic analyses. To aid forensic experts in their investigations, an image recognition model for microtrace localisation and classification is needed. In this work, we use deep learning to automate trace recognition in images captured with automated microscopy. We localise and classify fibres, hairs, skin, glass and sand in microscopy scans through pixel-wise classification of tape-lift samples. As deep learning requires extensive amounts of annotated training data, we additionally investigate various pretraining strategies to minimise the required annotation workload. We compare ImageNet pretraining, pretraining with self-supervised learning and a sequential application of these approaches. We find that pretrained models are able to reduce the required annotated data twofold compared to models trained from scratch while retaining the prediction accuracy. While our ImageNet-pretrained models outperform our self-supervised-pretrained models, we achieve the highest accuracy by combining the two approaches, resulting in a factor of 4 reduction of manual annotated microtraces or a 65 % improvement in recognition and localisation accuracy (mean intersection over union increases from 0.34 to 0.56 due to pretraining) when training on only 2.2 dm^2^ of annotated tape lift scans. Our model is, therefore, the method of choice for the automatic analysis of large forensic microtrace scans.

## 1. Introduction

Microtraces such as hairs, skin cells and fibres often aid in the reconstruction of a crime by providing information on items, locations, people and their actions [1]. Microtraces need to be recovered from their carrier to allow a detailed analysis. One of the used methods to recover these traces from areas of interest is tape lifting. Tape lifting is a fast technique and allows the recovery of different types of traces. [2]. A specialized method to lift traces from an item or body is one-to-one taping [3, 4]. In this method, the item or body is fully covered with transparent tapes, such that the original location of the traces identified on the tapes can be retrieved. This may provide important clues in activity level interpretations [5].

Tape lifting is a quick and efficient method to recover traces. In the next step, the recovered traces need to be examined [2, 6, 7]. Examination of traces on tapes is generally expensive, as it is a time-consuming task carried out by highly trained examiners. Automation of trace investigation will therefore reduce manual labour and cost and hence improve the effectiveness of trace evidence investigations. A recent step in automation has been made with the procurement of automated microscopy systems within the *Shuttle* project [8] to capture digital scans of tape-lifts. A key challenge that remains is the recognising and localising microtraces in the captured scans automatically and rapidly.

Therefore, we develop methodology for segmentation and classification in microscopic scans and show its ability to precisely localise and classify traces on tape-lifted microtrace samples. This model creates overviews that show the identity and distribution of some of the most useful microtraces. These overviews assist experts in deciding on further investigation procedures.

The developed methodology is based on a deep learning approach. Deep learning image recognition relies on training a neural network to derive meaningful information from input images.

Neural networks learn complex patterns and relationships within images. As a result, deep learning has achieved remarkable success in visual tasks such as image classification and semantic segmentation [9–11]. Typically, the network is trained with a large database of annotated images [10–14].

Our developed methodology includes a deep neural network for automated trace classification, schematically represented in Figure 1. In Figure 1a, we classify pixels in microscopy scans as either fibre, hair, skin, glass, sand or background. The network is composed of a ResNet-50 [10] feature extractor that embeds the relevant information of the input image into a feature representation. This feature representation is subsequently used by a fully convolutional [15] pixel classifier to assign a prediction for each pixel (semantic segmentation). The network is trained from scratch by initialising the weights with random parameters and training only via annotated microtrace images. Alternatively, we use an ImageNet-pretrained feature extractor (see Figure 1b) or pretrain the feature extractor with self-supervised learning (SSL) on unannotated microtrace images (see Figure 1c). In Figure 1d, we show our method for extracting and augmenting training images from microtrace scans.

**Figure 1:**
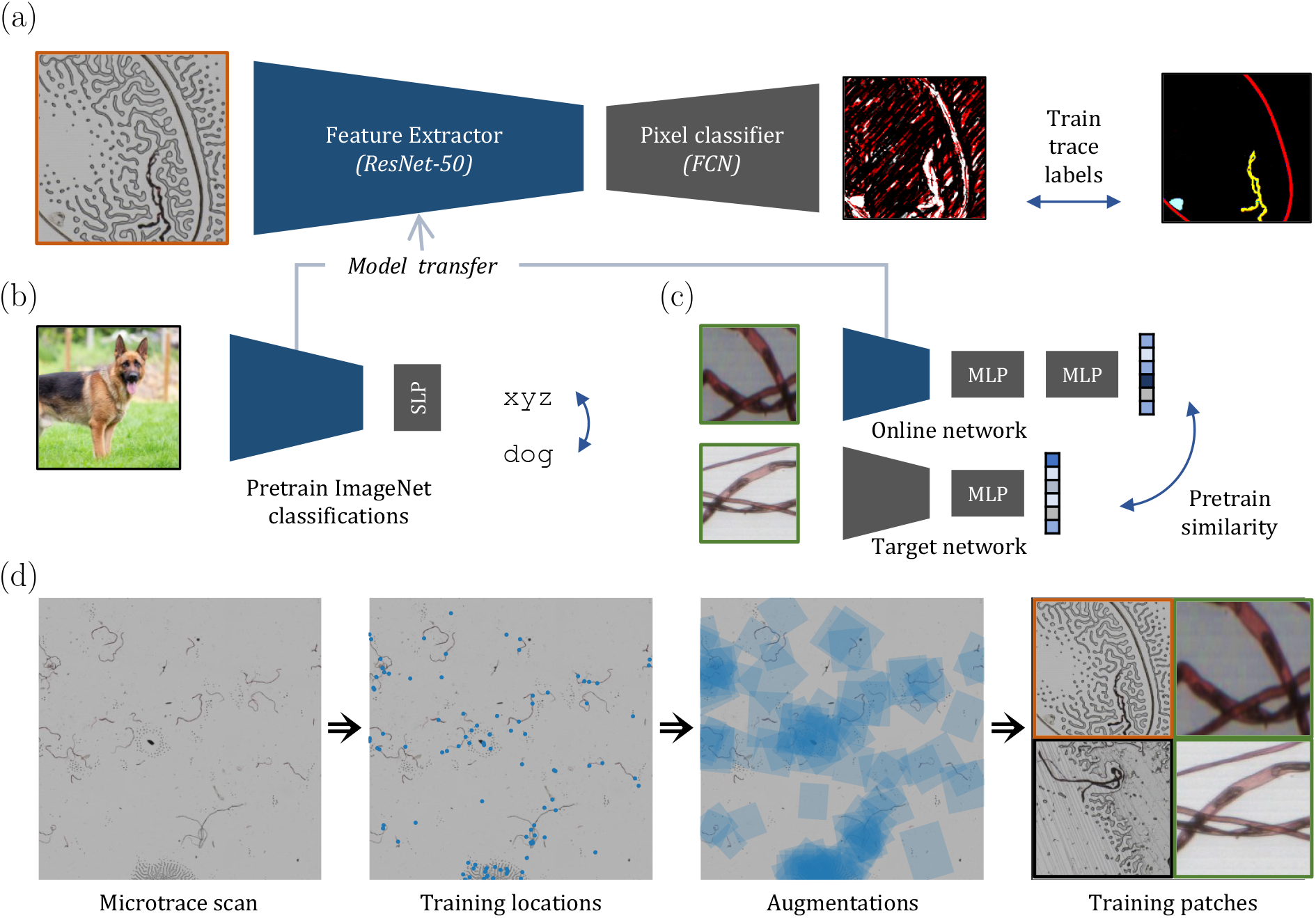
Schematic overview of deep neural network for automated microtrace classification. (a) Training trace predictions with expert annotations. A residual network with 50 layers (ResNet-50) [10] feature extractor is used in a fully convolutional network (FCN) [15] structure for pixel-wise classification. The network aims to find the relationship between the microtrace images (left) and the corresponding annotation provided by experts (right) by minimising the cross-entropy loss between the predictions and the annotations. The microtrace image in the figure shows a hair, a fibre and air bubbles. (b) ImageNet pretraining of feature extractor through classification of annotated everyday photographs in the ImageNet database [17] with a single layer perceptron (SLP) classifier. (c) Pretraining the feature extractor with the self-supervised learning (SSL) model Byol [22] on pairs of unannotated images. Here, the weights of the feature extractor are adapted such that pairs of similar images yield similar features. The architecture is composed of two ResNet-50 networks and a set of multi-layer perceptrons (MLPs). (d) Extraction and augmentation of training images from tape-lift scans. For pretraining with SSL, image pairs are extracted. For training trace predictions, image patches are extracted together with their corresponding annotation mask.

Generally, the quality of a deep learning model improves with a larger set of annotated images. As an example, ImageNet, a well-known database, contains over 1 million annotated images that can be used to train image recognition models. These images show everyday objects and cannot be used to directly train a system to recognise images of microtraces. As a result, the development of a trace recognition model requires the creation of a database of annotated microtrace images.

As an additional requirement, forensic experts are interested not only in the presence or absence of specific traces but also in their location or spatial distribution. Hence, the annotations should also provide the segmentation and location of the traces. This implies that each pixel in the training data has to be annotated. Altogether, a large number of tapes has to be annotated by experts, as a variety of traces can be encountered with tape lifts and the number of traces per sample is low [16]. This makes the compilation of an annotated database a labour-intensive process that needs to be carried out by experts who can recognise and localise the relevant traces. The cost to compile a database may thus become extremely high [11].

Therefore, we additionally investigate methods to minimise the annotation workload through pretraining. In a pretraining task, the neural network is trained either on relevant images from other sources, or on unannotated images. It is anticipated that pretraining reduces the number of annotated images needed for robust recognition of traces.

The first pretraining method used in the current study is based on ImageNet classification [17, 18]. As stated before, ImageNet cannot be used to train a network to recognise microtraces. Nevertheless, training using ImageNet teaches the network to recognise visual features important for human observers, such as the colour, shape, and texture of objects in the input images. As the network already recognises such important features, recognition of additional items, such as microtraces, may become easier. Indeed, it has been shown that a pretrained neural network outperforms a randomly initialised network with regard to convergence speed and accuracy [19]. Furthermore, ImageNet pretrained models are widely available, and no computational efforts are needed to use a pretraining neural network [20]. However, the benefits of ImageNet pretraining diminish for tasks dissimilar to the classification of everyday photographs, as the learned visual features can be sub-optimal for other domains [18, 21].

The second pretraining method is based on images of microtraces that have not been annotated. This approach is based on self-supervised learning (SSL) and uses the framework proposed in [22]. In this approach, two different and perturbed versions are made for each unannotated microtrace which are called *augmentations*. The two augmentations are made via cropping, recolouring and rotating. Pretraining aims to minimise the distance between the features of the two augmentations. In this way, the neural network learns what features of an image are important for microtrace classification and are to be used when microtraces are observed.

In SSL, the system is trained on augmented images, without knowing the nature of the displayed images. Therefore, pretraining by SSL does not require the annotation of images.

Finally, we combined the pretraining approaches: a neural network is pretrained using ImageNet pretraining. Subsequently, the network is further pretrained using SSL, to optimise the processing of microtraces. This approach follows recent research on pretraining in SSL [23].

## 2. Methods

### 2.1. Samples and datasets

Textile, glass and sand samples were taken from the general collection available in the authors’ laboratory. Dandruff, donated by volunteers was used as skin cells. The used tapes and tape backings were provided by Spectricon (Chania, Greece). Samples were prepared by distributing materials on a solid surface and lifted using the tape. Afterwards, the tape was attached to the provided backing to prevent contamination. Images of the tapes were acquired using a smmart automated microscope, developed by Spectricon, Chania, Greece [24], within the context of the shuttle tender [8]. Samples were imaged using transmitted light and acquired as colour images, using the calibration and acquisition procedures proposed by the manufacturer. Each pixel in the resulting images represents an area of approximately 1 *×* 1 μm. Images were corrected for shading effects and stitched together using routines written in Python. In this way, a tape of 80 *×* 80 mm results in a `.tiff` file that contains 6.4 Gigapixels. In total, 324 tape samples of various sizes were scanned. Ten of these were annotated (see Table 1).

**Table 1:** Overview of datasets. The last column shows the number of unique image patches that can be extracted for a field of view (FOV) of 1024 *×* 1024 μm. The unannotated dataset includes the tapes of the annotated training dataset. Thus, a total of 315 samples were scanned.

| Dataset | #tapes | Total area | #patches |
| --- | --- | --- | --- |
| Unannotated | 314 | $1.082 \text{ m}^2$ | $1.0 \cdot 10^6$ |
| Annotated |  |  |  |
| Training | 9 | $0.022 \text{ m}^2$ | $21 \cdot 10^3$ |
| Test | 1 | $0.006 \text{ m}^2$ | $5.8 \cdot 10^3$ |

### 2.2. Annotation

Traces present in a selection of the acquired images were segmented by thresholding and preliminarily classified using simple heuristics, such as the colour histograms and selected shape features. The preliminary annotations were saved as geojson objects and opened in QuPath software [25] together with the original images, where experts manually corrected the preliminary annotations. In this way, the workload for the experts was minimised. The time spent on correcting and manually annotating trace images is estimated around 120 hours. In this period, a dataset of approximately 21.000 annotated image patches (see Table 1) was created. Table 2 presents the occurrence of each trace annotation in the training dataset.

**Table 2:**
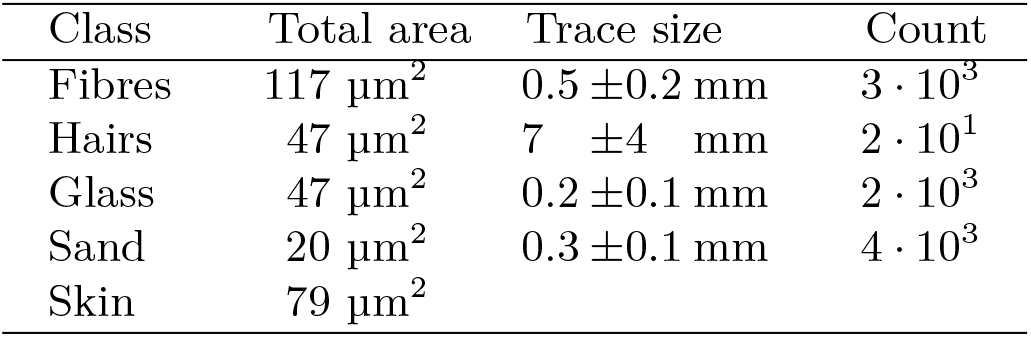
Traces in annotated training dataset. The class skin is segmented in regions instead of individual cells, therefore yielding the two right columns ill-defined. The trace size is calculated as the median diameter of the minimum bounding circle for each annotation. For the uncertainty measure, the median absolute deviation is used.

The annotated dataset was split into training and test sets. The test set consisted of an 80 *×* 80 mm tape lift containing all trace types. Nine other annotated tapes were used as training set. In total, the tape lift area of the training set was approximately three times as large as the test set, as shown in Table 1.

### 2.3. Training

A schematic representation of the neural network used for microtrace classification is provided in Figure 1a. A detailed overview of the architecture is provided in Supplementary Figure S7. This is based on the procedure proposed elsewhere [22, 26]. The network consists of a fully convolutional network (FCN) architecture [15] with a ResNet-50 feature extractor [10]. The last block of the feature extractor (*stage 5*) uses dilated convolutions to increase the resolution of the feature map, which may benefit semantic segmentation [27].

The network is trained by minimising the difference between the predicted pixel classifications and the expert annotated classifications via standard per-pixel softmax cross-entropy loss [20, 28]. We do not freeze the weights of pretrained feature extractors but instead allow all network weights to be optimised.

In benchmark tests, the feature extractor was randomly initialised. Supplementary Table S1, S2 and S3 respectively provide the hyperparameters, augmentations and initialisations used in training.

### 2.4. ImageNet pretraining

For the ImageNet pretraining approach, the neural network architecture and training procedure are identical to the procedure presented in Subsection 2.3. However, the ResNet-50 weights are not randomly initialised. Rather, they were initialised by the model provided by [20]. This procedure is visualised in Figure 1a and Figure 1b. Figure 1b shows the training of a model based on the 1.3 million images of ImageNet [17]. This training was carried out by [20]. The trained network was downloaded and its weights were used to initialise the network for training the classifier using microtrace images. This transfer is illustrated by the arrow between the subfigures of Figure 1. The hyperparameters used by [20] to obtain these weights are summarised in Supplementary Table S1.

### 2.5. Self-supervised learning

Figure 1c shows pretraining of the feature extractor using SSL on image patches extracted from unannotated microtrace scans. We employ the SSL model “Bootstrap Your Own Latent” (Byol) [22]. With Byol, the feature extractor is rewarded for predicting the similarity of the feature representations of two images with the same underlying structure.

These pairs of images are obtained by augmentation, i.e. alteration of an image of a single trace. For this purpose, an image is retrieved using the method described in Subsection 2.6. This image is augmented twice by using a combination of translation and zoom parameters. Cropping is an essential aspect of creating useful image pairs [22]. Cropping is performed by adjusting translation and zoom parameters, which were consequently optimized for better results. For further details, refer to Supplementary Figure S4.

Byol is composed of an asymmetric architecture of two separate networks: an online network and a target network, as shown in Figure 1b. The online network is composed of a ResNet-50 feature extractor, an multilayer perceptron (MLP) projector and an MLP predictor. The target network is composed of a second feature extractor and a second projector, both with different weights than the online network. Supplementary Figure S8 provides a detailed overview of the implementation and the output dimensions of each network.

During SSL, the online network and target network each take one of the augmented patches as input. The weights of the online network are optimised to predict the output of the target network.

In this way, the model learns to recognise the similarities between two cropped parts of a single trace image and, hence, to recognise the important features on which the recognition can be based. SSL does not require a classification annotation of the trace. As a result, it can learn from images that have not been annotated. The implementation of SSL is discussed in detail in Supplementary Note 1.

### 2.6. Image Extraction

The training (see Subsection 2.3) and SSL (see Subsection 2.5) make use of image patches. These image patches are extracted from transmission microscopy scans as shown in Figure 1d.

Tape lifts generally contain a variety of small traces scattered on a transparent tape. The dimensions of the used tapes are 80 × 80 mm, which is very large compared to the size of common microtraces and to the optical resolution of the used microscopy system. This results in large datasets containing around 6.4 Mpixel. The tapes are usually not completely covered by microtraces, but also contain background areas. In fact, experts annotated less than 2% of all pixels as trace (see Supplementary Table S4). The large size of these scans prevents calculations on a full scan, as the required computing power would be immense. Instead, image patches are sampled from the scans.

Random sampling image patches from the microtrace samples [11] would result in many patches containing only background, which is not useful for training.

Fortunately, most of the background area can easily be recognised. The used tape has an excellent light transmission and the background is shown as homogeneous white areas. Traces generally scatter and absorb light and are thus displayed as dark objects. Therefore, thresholding with a single global threshold suffices to distinguish foreground areas from background [29]. The threshold value is determined using the histogram-based triangle method [30], which is suited for images dominantly consisting of background [29, 31, 32]. This is shown in Supplementary Figure S11. With the found threshold, a total of 94% of the image area in the annotated dataset is estimated as background. The remaining 6% is estimated as foreground, including possible microtraces or artefacts. The proposed procedure extracts patches centred around foreground areas. As a result, filtering out background-rich areas later in the process, such as proposed by [19, 33] is not necessary. Training coordinates are then sampled uniformly at random from the thresholded foreground areas. At each sampled coordinate, an image patch of size 256*×*256 pixels is extracted in a resolution of 4 μm per pixel.

For trace training, single image patches are extracted together with corresponding annotation masks. These image patches are augmented to artificially enlarge the dataset. Specifically, we make a crop of random zoom, aspect ratio and rotation and resize it to 256 *×* 256 pixels as shown in Supplementary Figure S12. Then, we further augment the image patches by recolouring, blurring and mirroring, which results in the required training images. The chosen augmentations, randomisation processes and their parameters are derived from [22] and are listed in Supplementary Table S2.

The chosen field of view (FOV) of 1024 *×* 1024 μm per patch ensures that the majority of the traces is captured fully within the FOV (see Table 2). Images are mean-downsampled to a resolution of 4 μm/pixel, resulting in a patch size of 256 *×* 256 pixels.

### 2.7. Evaluation

Trained models are tested by evaluating their predictions on an unseen microtrace scan. The used scan was annotated, but was not part of the annotated training dataset or the unannotated dataset used in SSL pretraining. This scan consists of an 80 *×* 80 mm tape lift sample containing all trace types, shown in Supplementary Figure S14. During testing, the scan is processed via a uniform grid of image patches (see Supplementary Figure S5), without oversampling the foreground with thresholding or applying augmentations as described in Subsection 2.6. For each image patch, the predictions at the edges are discarded, following [28, 34], due to their lower accuracy. The FOV of each image patch is increased during testing with 12.5%, resulting in partially overlapping input patches.

The trace recognition and localisation performance of the model is evaluated with the Intersection over Union (IoU) [11, 35]. For any class *A*, the IoU is defined as the number of correctly identified trace pixels divided by the sum of all pixels either predicted or annotated as class *A*:

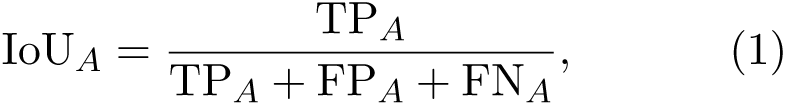

Here, TP_*A*_ denotes the number of true positives, while FP_*A*_ and FN_*A*_ denote the number of false positives and false negatives, respectively. The maximum value of IoU_*A*_ = 1 indicates that the annotations and predictions match perfectly. Conversely, the minimum score of IoU_*A*_ = 0 indicates no true positives were predicted.

To quantify the overall performance over all classes, the mean of the IoU (mIoU) of the *n*_*c*_ non-background classes is taken:

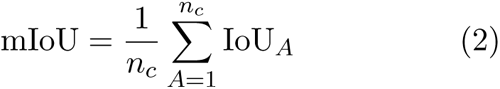

In our case, *n*_*c*_ = 5, with the classes being fibre, glass, hair, sand, and skin. Similar to the evaluation protocol of [12], we calculate the overall mean Intersection over Union (mIoU) based on the accumulated statistics over the entire test set. As single image patches typically do not contain all of the five trace classes, the accumulated statistics are a more generalizable representation of the trace recognition performance. The violin plots used throughout the report show the distribution of mIoU values that are obtained by accumulating over a randomly determined 10-fold split in the test set that was kept constant throughout the experiments.

mIoU is calculated by averaging over the classes present in the test sets. This means that the IoU of each class contributes equally to the mean regardless of how many examples of each class are present in the test dataset. This approach is also known as micro-averaging [36], in contrast to macro-averaging, which aggregates the IoU scores across all instances in the test set. Macro-averaged IoU can be more lenient towards biased classifiers in imbalanced datasets due to its aggregation of class-wise IoU values. Therefore, it is advised to employ a micro-averaging approach for unbalanced datasets. Moreover, we display violin plots with the mIoU distributions and show the confusion matrices (Supplementary Figure S1) to provide additional insight into the model performance.

To investigate annotation-efficient learning, we split our annotated training set into equally sized regions as shown in Supplementary Figure S15 and vary the number of regions used in training the trace classifications.

## 3. Results

Four models were trained, following the general procedure in Figure 1, namely models with random initialisation, with ImageNet pretraining, with SSL pretraining, and with the combination of both pretraining methods. Our results show that the final option, combining both pretraining methods, achieves the best classification, as indicated by the highest mIoU. This training will be highlighted in Subsection 3.1. Subsection 3.2 presents our results on the benefit of pretraining, followed by a discussion of our results on SSL in Subsection 3.3. Finally, we provide insight into the model and its key elements through an ablation study, that is by removing parts of the pipeline. We discuss the results obtained by these ablated methods in Subsection 3.4.

### 3.1. Microtrace recognition

The model described in the current section was initialised by an ImageNet pretrained model. It was further pretrained using SSL on roughly 1 m^2^ of unannotated tape area. Finally, the model was trained on our annotated dataset of 22 dm^2^ of tape area. This training method resulted in an mIoU of 0.56.

A general impression of the meaning of the value of mIoU can be obtained from Figure 2. This figure provides three rows of images. The top row provides images as acquired by the automated microscope. These images were part of the test set, so they have been annotated by an expert, but were not used during the training of the model. Rather, the trained model was used to identify and localise traces. As a result, the annotations and predictions can be compared. The second row of images provides the annotations. These annotations are colour-coded using the colours shown below the figure. The third row of images provides the predictions generated by the model, using the same colour-coding. Column a shows an image of air bubbles and other artefacts without annotated traces. Here, the model aligns with the expert in not marking any traces. Column b shows a fibre. Here, the model classified part of the trace as fibre and incorrectly masked the largest part of the trace as hair. This leads to a fibre mIoU of 0.28 and a hair mIoU of 0.00. Column c and d show correctly identified hairs. Due to imperfect masks, IoUs of 0.77 and 0.76 are attained respectively. Column e shows a trace that was classified as sand by both the model, and the expert. The areas indicated by the model and the expert overlap almost perfectly, leading to an IoU of 0.95. Column g shows an area in which glass particles are annotated and predicted. A visual comparison of these images indicates that all glass particles were found by both the expert and the model. Nevertheless, the IoU is only 0.72, indicating that the areas shown do not perfectly overlap. In column h, the situation is even more extreme. The model predicts the same as the expert annotated. Yet the calculated IoU is only 0.24, as the model missed part of the fibre.

**Figure 2:**
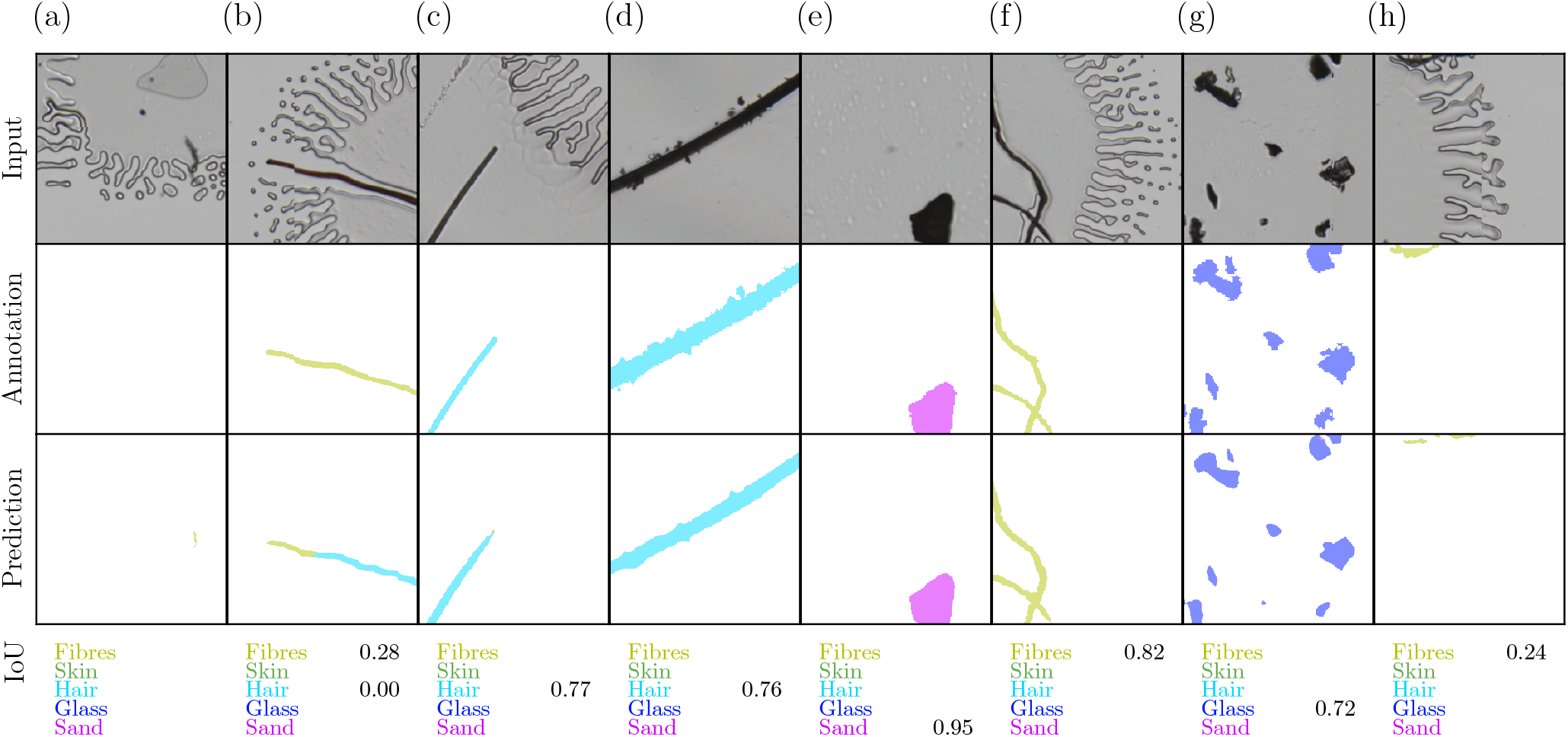
Comparison of model predictions with expert annotations for patches in the test set to visually assess model performance and corresponding IoU values. The IoU values for classes that span less than 1% of the total image area in both the expert annotations and the predictions are omitted.

It should be noted that the mIoU is based on the number of correctly classified pixels. Forensic examiners are usually more interested in the number of identified microtraces. As each of the microtraces comprises several pixels in the images, the number of traces found is higher than the mIoU. In practice, we find that nearly all traces are located, though our current methodology does not allow an accurate determination of the number of accurately predicted traces.

Supplementary Figure S2c provides additional image patches on which the model performs poorly. These images were randomly selected from predictions where a class IoU of less than 0.4 was encountered.

Air bubble patterns can be seen in most of the microscopic images. These are caused by air being trapped under the tape. It can be seen from Figure 2 that the model correctly refrained from identifying these patterns as traces.

Supplementary Figure S1 shows the confusion matrix of the model together with precision and recall analysis. It can be seen that the pixel-wise precision exceeds 60% for each class, indicating that at least 60% of the pixels marked as a certain class are marked as that class by the expert annotator as well. Except for the class Skin, the recall exceeds 60% as well, indicating that at least 60% of the pixels are marked as a certain class by both the expert and the model.

Generating predictions for a microtrace tape of 80 *×* 80 mm takes approximately 30 seconds using our hardware (nvidia rtx3090 GPU and Intel i910900X CPU). This is a fraction of the time required for a microscopist to generate an overview of traces on the tape. Also, it is much faster than scanning the tape with automated microscopy, which takes around 40 minutes with the used device.

### 3.2. Pretraining for annotation-efficient learning

The current section explores whether pretraining reduces the time-intensive and hence expensive task of annotating images. Our main results are presented in Figure 3. The horizontal axis in this figure represents the area of tape that was used to train the network. Note that this axis is not linear, as it is based on a sequential doubling of the number of tapes used for annotation. The vertical axis shows the accuracy of the trained model, as indicated by the mIoU. Training on smaller areas reduces the diversity of the microtraces used to train the model. Therefore, the predictions become less accurate. Therefore, the numbers presented in Figure 3 for areas of 0.1, 0.3, and 0.6 dm^2^ are averaged over 5, 3, and 2 training runs, respectively.

**Figure 3:**
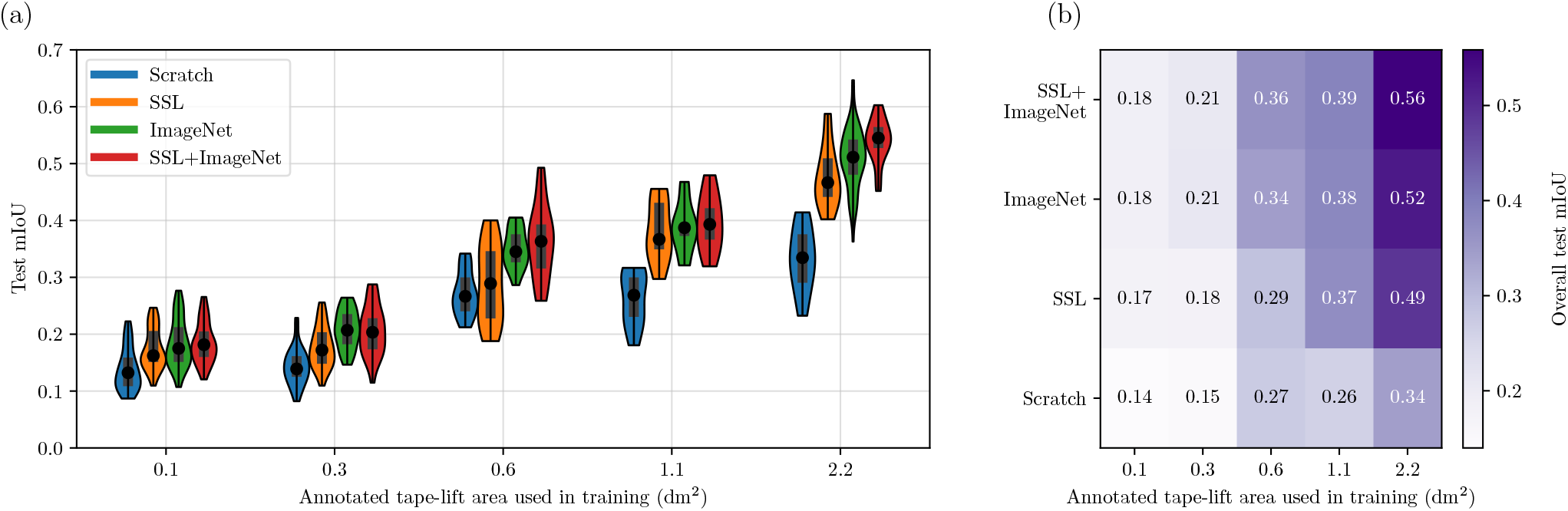
Benefit of pretraining for annotation-efficient learning. (a) Violin plot showing the distribution of mIoU over a 10-fold split in the test set. The dot presents the median, the inner bar the upper and lower quartile. The line presents the spread between the minimum and maximum. (b) Overall test mIoU per experiment.

Figure 3 shows that training on a larger area improves the accuracy of prediction. In addition, the pretrained models achieve higher mIoU values than the models trained from scratch. The ImageNet-pretrained models outperform the SSL-pretrained models. The highest mIoU value is obtained for the model pretrained by both ImageNet pretraining and SSL, followed by training on 2.2 dm^2^.

When the model is trained on 0.1dm^2^, pretraining improves the mIoU from 0.14 to 0.18, which is an increase of 29%. The effects of pretraining become even higher when the model is trained on larger areas. When the model is trained on 2.2 dm^2^, pretraining improves the mIoU from 0.34 to 0.56 for the combined pretraining methods, i.e. an improvement of 65%. In addition, pretraining followed by training on 0.6 dm^2^ of annotated tape or more leads to models that outperform models trained from scratch using two times more annotated data. In fact, combined ImageNet and SSL pretraining followed by training on 0.6 dm^2^ of annotated tape leads to a higher mIoU (0.36) than a training from scratch using 2.2 dm^2^ (0.34). This means that pretraining, in this specific case, reduced the annotation workload by a factor of four.

Values of mIoU below unity indicate that the classification achieved by the model is not perfect. The types of errors made by the models are not specified in Figure 3, but can be identified by a further exploration of the data. Supplementary Figure S3 shows a detailed exploration of the various error modes for our experiments. When the model is trained on 2.2 dm^2^, the 65% improved mIoU after pretraining can be traced back to a 40% increase in correctly classified trace pixels, a 27% decrease in missed trace pixels and a 56% decrease in false detections compared to training from scratch. The number of confused trace pixels is increased by 1% after pretraining.

### 3.3. Self-supervised pretraining

SSL pretraining minimises the distance between the two different images or image augmentations of a similar trace. In the current study, pairs of image augmentations were obtained by selecting a coordinate on a microtrace scan, extracting two images within a maximum distance *d*_*m*_ of this point. Next, each of these images is randomly augmented, according to the procedure described in Subsection 2.6). Combinations of several augmentation methods have been tested. An overview is presented in Supplementary Table S2.

The results of the SSL experiments of Subsection 3.2 were obtained with *d*_*m*_ = 0 and a maximum magnification augmentation between 0.5× and 2×. These values lead to optimal results, as shown in Supplementary Figure S4a, which shows the obtained mIoU values for different settings of the magnification and translation. An mIoU of 0.42 is achieved with *d*_*m*_ = 0 and magnification augmentation between 0.5× and 2×, while choosing a lighter zoom augmentation (between 0.67× and 1.5×), results in an mIoU of only 0.37. It can be anticipated that a light zoom augmentation and no translation results in two approximately similar representations for both views and hence a trivial solution. This reduces the benefit of pretraining.

Optimal parameters for the augmentation are dependent on the class of the trace involved. As stated before, SSL is not aware of the class of the trace involved. Nevertheless, it is possible to pretrain the model using different types of traces. Supplementary Figure S4b-f show the IoU benefit per class to illustrate this effect. This can be illustrated by the results for hairs and glass. A translation of 2048 μm hardly affects the IoU for hair traces (0.61 vs 0.62, see Figure S4d), while such a large translation reduces the IoU for glass traces from 0.48 to 0.34 (see Figure S4e). This is attributed to the size of the traces. Glass fragments are generally small. Table 2 shows that the glass particles of our dataset generally have sizes of 0.2−0.4 mm. A translation of 2048 μm can therefore cause the trace to move out of the field of view for one of the images in the augmentation pair. This obviously hinders the training, as the SSL procedure assumes that both images display items that should be considered similar. On the other hand, hairs have lengths of more than 5 mm, yielding a larger probability of encountering the trace in both views. The presented parameters are a compromise that offers a balanced performance for all trace types included.

### 3.4. Influence of Image Sampling

In the proposed method, training and SSL procedures make use of image patches extracted from transmission microscopy scans. The proposed method to extract the image patches, as described in Subsection 2.7 involves a thresholding operation, intended to enhance the amount of foreground pixels in the used image patches. Thresholding was considered beneficial due to the large amount of background in the investigated microscopic scans. In the current section, our method is compared to an alternative method that does not involve thresholding. Rather, this alternative method uses a uniform grid as shown in Supplementary Figure S5.

The results of this comparison are presented inFigure 4. The horizontal axis in Figure 4 shows the number of image patches used to train the model. The blue datapoints show the calculated mIoU when using the thresholding method. The performance of the model increases linearly with the number of images used for training, until it levels at an mIoU of around 0.55. If a uniform grid is used, (‘without thresholding’), a lower test mIoU was attained for each of the tested number of training images. After training on 400.000 image patches, the thresholding approach results in a 21% higher overall test mIoU compared to training with a uniform grid. Training on lower amounts of images further increases the performance difference. Specifically, when training with only 80.000 data points, our thresholding approach results in a four-fold increase in mIoU.

**Figure 4:**
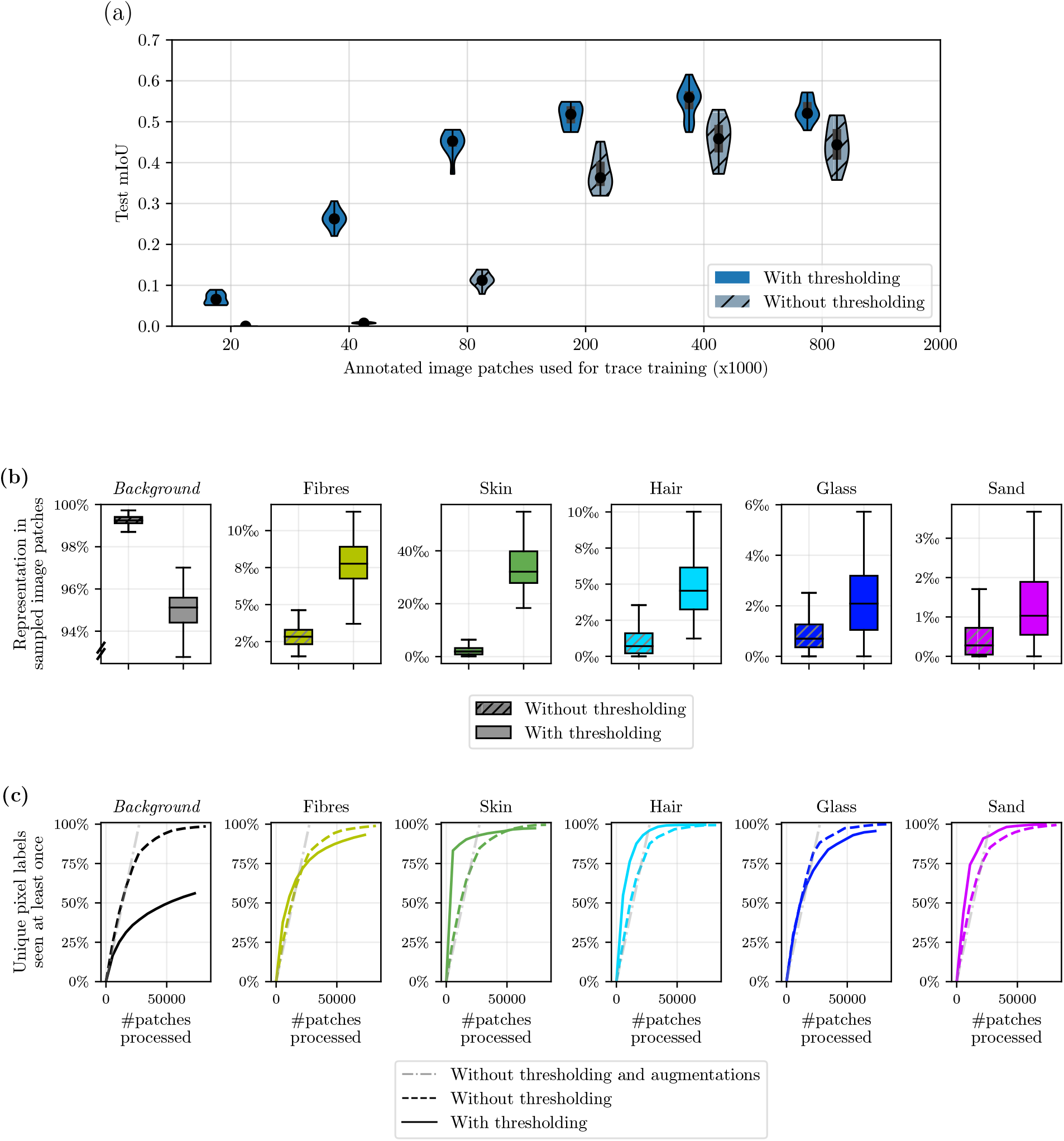
Benefit of thresholding approach shown in Figure 1d compared to the uniform approach shown in Supplementary Figure S5. Only trace training is regarded. An ImageNet pretrained-model is used without additional self-supervised pretraining. For the uniform approach, processing 21.000 image patches corresponds to one data set iteration (see Table 1). This is called one *epoch*. (a) mIoU of trace recognition for training with and without thresholding approach. The test mIoU is shown for training with various amounts of image patches across the annotated dataset. It can be seen that the thresholding approach converges earlier and to a higher final accuracy. (b) Efficacy of thresholding approach to oversample the foreground pixels. The class distribution of pixels over 8000 randomly selected image patches is shown. The left boxplots show the distribution of patches sampled without thresholding, the right boxplots show the class distribution for sampling with thresholding. It can be seen that the thresholding approach results in oversampling the non-background classes. (c) Analysis of missed pixels in the annotated dataset due to thresholding. It can be seen that after processing 80.000 image patches (4 epochs for the uniform approach), 95% of the non-background pixels have been seen at least once. Further experiments show that after 400.000 image patches (20 epochs for the uniform approach), 99.7% of all non-background pixels are seen, while 80% of the unique background pixels are seen.

The improved performance is attributed to a higher representation of traces in the used image patches. Further tests, detailed in Figure 4b and c, confirm the expectation that thresholding increases the representation of traces.

### 3.5. Influence of hyperparameters

The current section investigates the hyperparameters for training trace classifications. The sensitivity of segmentation accuracy with respect to hyperparameters is investigated and the chosen values are substantiated. These results were obtained after ImageNet pretraining. S6a shows that the mIoU of trace recognition has values between 0.5 and 0.6 for weight decay values between 0 and 10^*−*4^ but decreases below 0.05 for a decay value of 10^*−*2^. S6b furthermore shows that mIoU does not significantly change across batch sizes between 5 and 200 when the learning rate is scaled linearly with the batch size.

Images of traces contain many pixels. Each of these pixels is an individual input for our data model. Hence, the time needed to train a model increases with the number of input pixels. Downsampling of the microtrace images may help to limit the calculation times and reduce excess details that can lead to overfitting. On the other hand, downsampling images reduces the details that may be important for classifying traces. Therefore, we evaluated the performance and calculation times of models trained on the original images (1 μm/pixel) and downsampled images (2 and 4 μm/pixel). The results are shown in Figure S6c and d. Figure S6d shows that downsampling to 2 μm/pixel results in a shorter training time (approximately 4 instead of 15 hours) and Figure S6d shows an unaffected high mIoU (0.6 mIoU). This results in a speed-up of 3.75 times.

The image patches extracted from the microtrace scans can be altered before they are used to train the model. Such alterations may improve the robustness of the training, as reliance on feature diversity in the training data is reduced. Hence, they may improve the quality of the classification [37]. However, excess alteration can cause the simulated feature diversity to transcend the feature diversity of the actual traces, causing a decay in segmentation accuracy. For example, heavily shearing an image of a trace can cause the image to not resemble the trace anymore. S6e and f show that altering the magnification and aspect ratio has only a limited effect on the obtained mIoU values. Changing the magnification improves the model (mIoU 0.55 for magnification 0.33*×* and 3*×*, mIoU 0.52 for a magnification of 1. A limited alteration of the aspect ratio hardly influences the obtained mIoU values. Heavier zoom and aspect ratio augmentation deteriorate the training process. This is probably caused by the induced loss of information by these heavy augmentations.

## 4. Discussion and Conclusion

A data model is presented to classify microtraces shown in microscopic images. The model is based on a residual network with 50 layers (ResNet50). The presented model successfully localises and classifies hairs, fibres, skin and glass traces in the presented, resulting in a mIoU of 0.56. Classification of a data set representing a tape of 80 *×* 80 mm requires approximately 30 seconds.

The model was trained using images annotated by experts. As annotating is a time-consuming task, we explored pretraining methods to minimise the time needed for annotating. A pretrained ImageNet model can be downloaded, so its use does not incur any computational or annotation costs. Pretraining using SSL does not require annotated images, but is demanding computationally. The pretraining approaches result in an mIoU of 0.52 and 0.49 respectively, yielding a +53% and +44% improvement with respect to training from scratch using the same number of annotations. Combining ImageNet pretraining and SSL is more effective than the use of the individual methods and results in the optimal mIoU.

The presented mIoU of 0.56 is based on a model that was pretrained and subsequently trained on annotated data. A model trained from scratch on the same amount of annotated data achieves an mIoU of 0.34. Our tests indicated that pretraining may reduce the required amount of annotated data by approximately 4.

The finding that ImageNet pretraining outperforms sole SSL pretraining is in line with [33], in which SSL pretraining is investigated for semantic segmentation of biomedical microscopy images, although here another SSL framework (SimCLR [38]) is used. Moreover, it aligns with [39], in which Byol is compared to ImageNet pretraining for semantic segmentation of natural images [39]. However, [22] and [40, 41], find that Byol and SimCLR result in higher accuracies than ImageNet pretraining for semantic segmentation of natural images.

SSL is based on pairs of images displaying the same type of trace. In the current study, such pairs were created by extracting different parts from a single image patch of a trace. The image patches have been distorted to improve the robustness of the pretraining. The parameters selected to extract and distort the image patches have been optimised. It is shown that the optimal values depend on the size of the displayed trace. The chosen values are a magnification between 0.5-2*×* and a translation of 0 μm.

As tape lift scans typically contain a small number of traces scattered across a large background area, 99% of the image area of our dataset represents background (see Table S4). Our proposed image extraction method involving thresholding allows a +0.10 mIoU increase with respect to processing the image area uniformly by effectively oversampling foreground areas.

The presented model exports predictions in the geojson format. The predictions can be displayed along with the original images using QuPath [25] software, which experts have also used to annotate images. This provides a user-friendly method to evaluate the predictions by the model. In this way, the results of the model can be used to initiate further investigations.

The presented model successfully recognises hairs, fibres, skin, glass and sand particles. These are important traces in many forensic cases. Future studies should aim to improve classification by including more types of traces, such as blood traces and pollen. In addition, we aim to improve subclassification, so that different kinds of fibres, hairs, and crystals can be classified. These additional classification tasks may benefit from alternative microscopy modes, such as reflection imaging to improve the classification of opaque, thick traces (see Subsection 3.1).

Our study sets a new baseline for forensic microtrace recognition and gives practitioners insight into the benefits of pretraining, the required annotation workload, augmentation parameters and efficient image extraction for microtrace recognition in tape lift scans. We provide trained models and facilitate analysis of model predictions through a graphical user interface [25] to yield a valuable tool for microtrace recognition that aids forensic experts in their investigation. Hereby, we contribute to automating the microtrace finding process and decreasing the human labour required for trace investigations.

## 5. Acknowledgements

We thank the Hellenic Police and Staffordshire University for sharing scan records of tape lifts and thereby contributing to the dataset. Furthermore, we gratefully acknowledge the financial support of the Netherlands Forensic Institute (NFI). Lastly, we acknowledge Spectricon for the support regarding the smmart automated microscope.

## Notes

### Competing Interest Statement

Carlas Smith reports administrative support was provided by Delft University of Technology Faculty of Mechanical Engineering. The other authors declare that they have no known competing financial interests or personal relationships that could have appeared to influence the work reported in this paper.

