## Supplementary Materials for "Reducing Manual Labour in Forensic Microtrace Recognition with Deep Learning"

### List of Supplementary Notes

S1 Details on Self-Supervised Pretraining . . . . . . . . 20

### Supplementary Note 1: Details on Self-Supervised Pretraining

BYOL [22] is composed of two networks; an online network and a target network. By introducing asymmetry and only optimising the online network, BYOL prevents the feature extractor from learning a collapsed solution [44]. In a collapsed solution, the model outputs a constant feature representation independent of the input image. This constant representation results in a perfect similarity for each image pair, while it is not useful for classification. Although the weights of the target network feature extractor and projector can be identical to the corresponding weights of the online network [45], BYOL updates the target network with an exponential moving average of the weights of the online network. Hereby, it provides a smoother target representation to optimise the online network on. This results in improved performance with respect to using identical weights [22].

To train the online network, the loss function $\mathcal{L}$ is calculated for pair of augmented images $(V_i, V_i')$ as

$$\begin{aligned}\mathcal{L} := &\left\| \frac{z_\theta\left(V_i\right)}{\left\|z_\theta\left(V_i\right)\right\|_2} - \frac{f_\xi\left(V_i'\right)}{\left\|f_\xi\left(V_i'\right)\right\|_2} \right\|_2^2 + \\ &\left\| \frac{z_\theta\left(V_i'\right)}{\left\|z_\theta\left(V_i'\right)\right\|_2} - \frac{f_\xi\left(V_i\right)}{\left\|f_\xi\left(V_i\right)\right\|_2} \right\|_2^2 .\end{aligned} \tag{3}$$

Here, the output of the online network $z_\theta(\cdots)$ is normalised and compared to the normalised output of the target network $f_\xi(\cdots)$. The sum of the mean squared error is taken for feeding $V_i$ to the online network and $V_i'$ to the target network and the other way around.

At each optimisation step $k$, the online network is optimised to minimise $\mathcal{L}$ and the target network receives an exponential moving average of the parameters of the online network $\theta$ (predictor excluded):

$$\theta_{k+1} = \texttt{optimiser}(\theta_k, \nabla_\theta \mathcal{L}, \eta) \tag{4}$$

$$\xi_{k+1} = \tau_k \; \xi_k + (1 - \tau_k)\, \theta_{k+1}. \tag{5}$$

We use a batch size $m = 80$ and thus process 80 image pairs per optimisation step. Following [22] for small batch sizes, the base learning rate is calculated as $\eta_{\text{base}} = 0.4 \cdot m/256$. The learning rate is scheduled to increase linearly (*warm-up*) from 0 to $\eta_{\text{base}}$ within 1250 steps (100.000 image pairs) and to decay to 0 at the maximum number of steps $K$ with cosine decay [46].

The weight decay is set to $1.5 \cdot 10^{-6}$ independent of the batch size $m$, following [22].

As proposed in [22], we schedule the exponential moving average parameter $\tau_k$ to increase to reach 1 at the maximum number of steps $K$ with

$$\tau_k = 1 - (1 - \tau_0)\; \frac{\cos\left(\pi k/K\right) + 1}{2}. \tag{6}$$

In [22], $\tau_0 = 0.996$ and $\tau_0 = 0.9995$ are proposed for batch sizes $m = 4096$ and $m = 512$ respectively. We further scale $\tau_0$ from a reference batch size $m_{\text{ref}}$ to a target batch size $m_{\text{tgt}}$ with the exponential moving average parameter scaling rule proposed in [47]:

$$\tau_{0,\text{tgt}} = \tau_{0,\text{ref}}^{m_{\text{tgt}}/m_{\text{ref}}}. \tag{7}$$

Filling in $\tau_{0,\text{ref}} = 0.996$, $m_{\text{ref}} = 4096$ gives $\tau_{0,\text{tgt}} = 0.9995$ for $m_{\text{ref}} = 512$, matching $\tau_0$ proposed in [22]. For our batch size $m = 80$, the scaling yields $\tau_0 = 0.99992$, which is the value we use in our experiments.

We pretrain with SGD [20] using a momentum factor of 0.9 on 10 million image pairs ($K = 125.000$) across the unannotated dataset. This costs four days of training on our hardware. In Supplementary Figure S10, we show test mIoU results obtained for training with fewer image pairs.

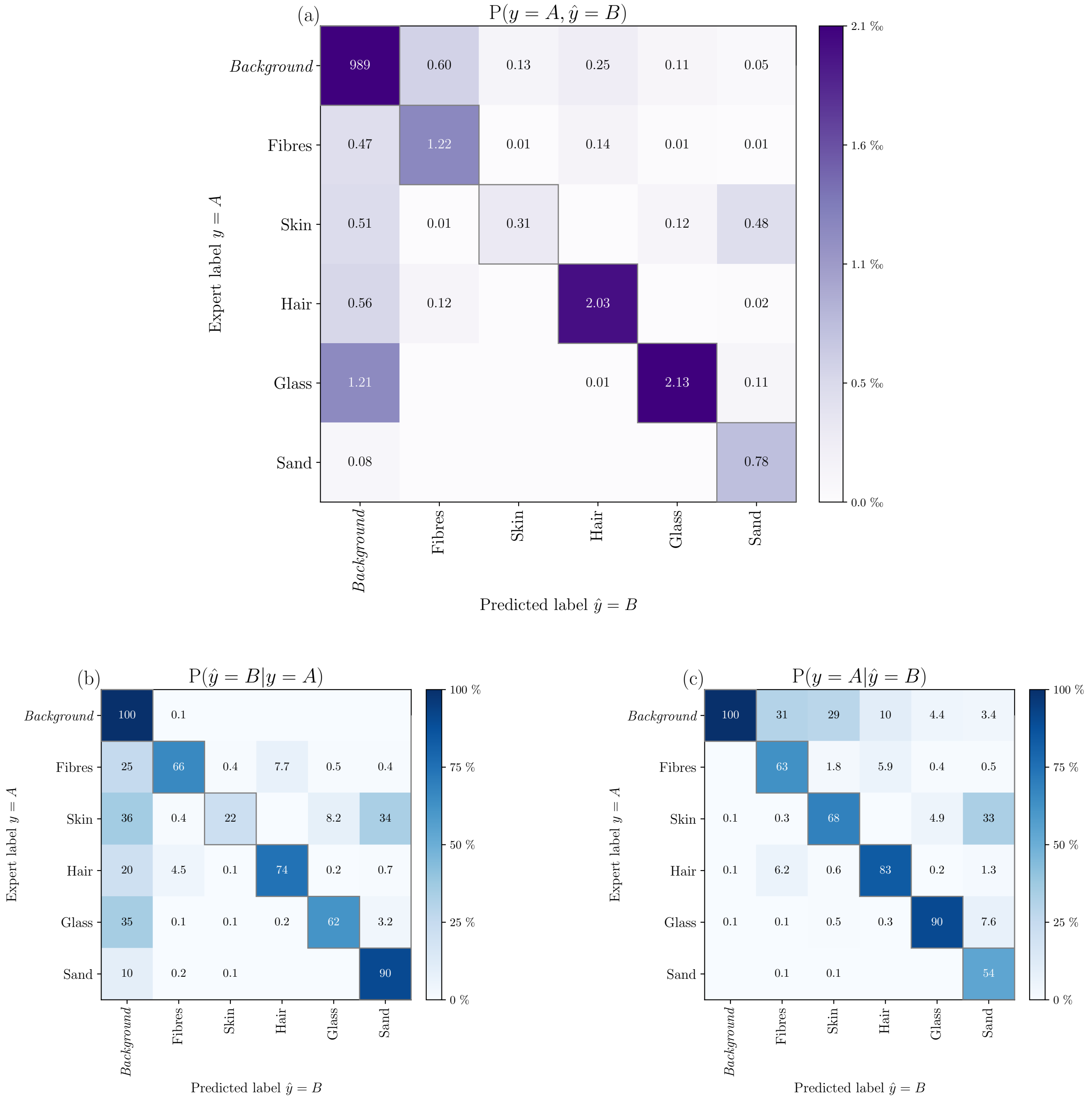

**Supplementary Figure S1:** Confusion matrix of predictions in test set. (a) Normalised with total number of predictions. Values lower than 0.005 ‰ are omitted. (b) Row-wise normalised, yielding probabilities conditioned with true values. The diagonal presents the recall (sensitivity) per trace class. Values lower than 0.05% are omitted. (c) Column-wise normalised, yielding probabilities conditioned with predicted values. The diagonal presents the precision per trace class. Values lower than 0.05% are omitted.

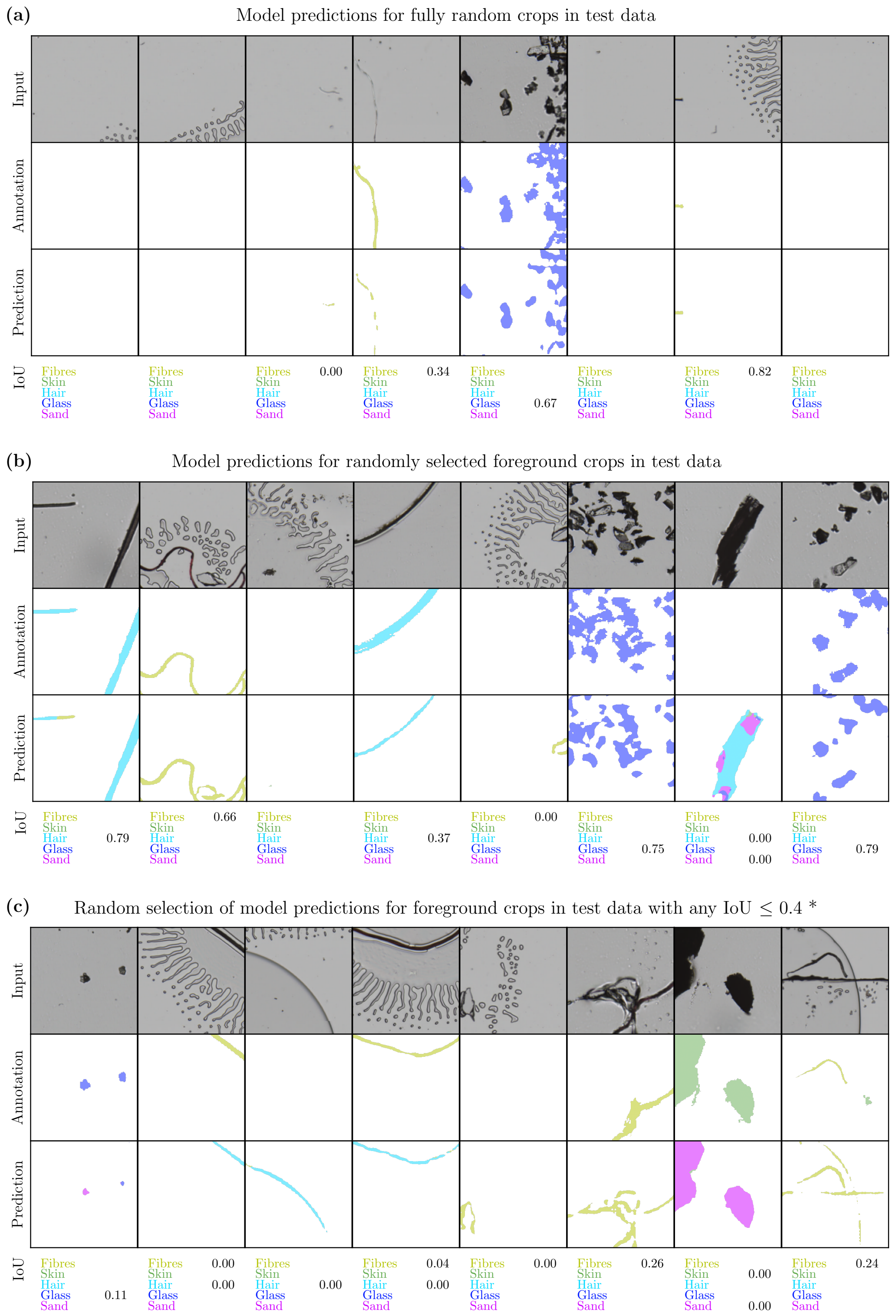

**Supplementary Figure S2:** Selection of model predictions in test dataset to visually assess model performance and corresponding IoU values. The IoU values for classes that span less than 1% of the total image area in both the expert annotations and the predictions are omitted. (a) Predictions for fully random crops in test data. Here, the location of the crops was selected via uniform sampling. (b) Predictions for randomly selected foreground crops. These crops were extracted with the thresholding approach discussed in Subsection 2.6. (c) Foreground crops where any of the IoUs for the classes that span at least 1% of the image area in either the predictions of the annotations, is lower than 0.4.

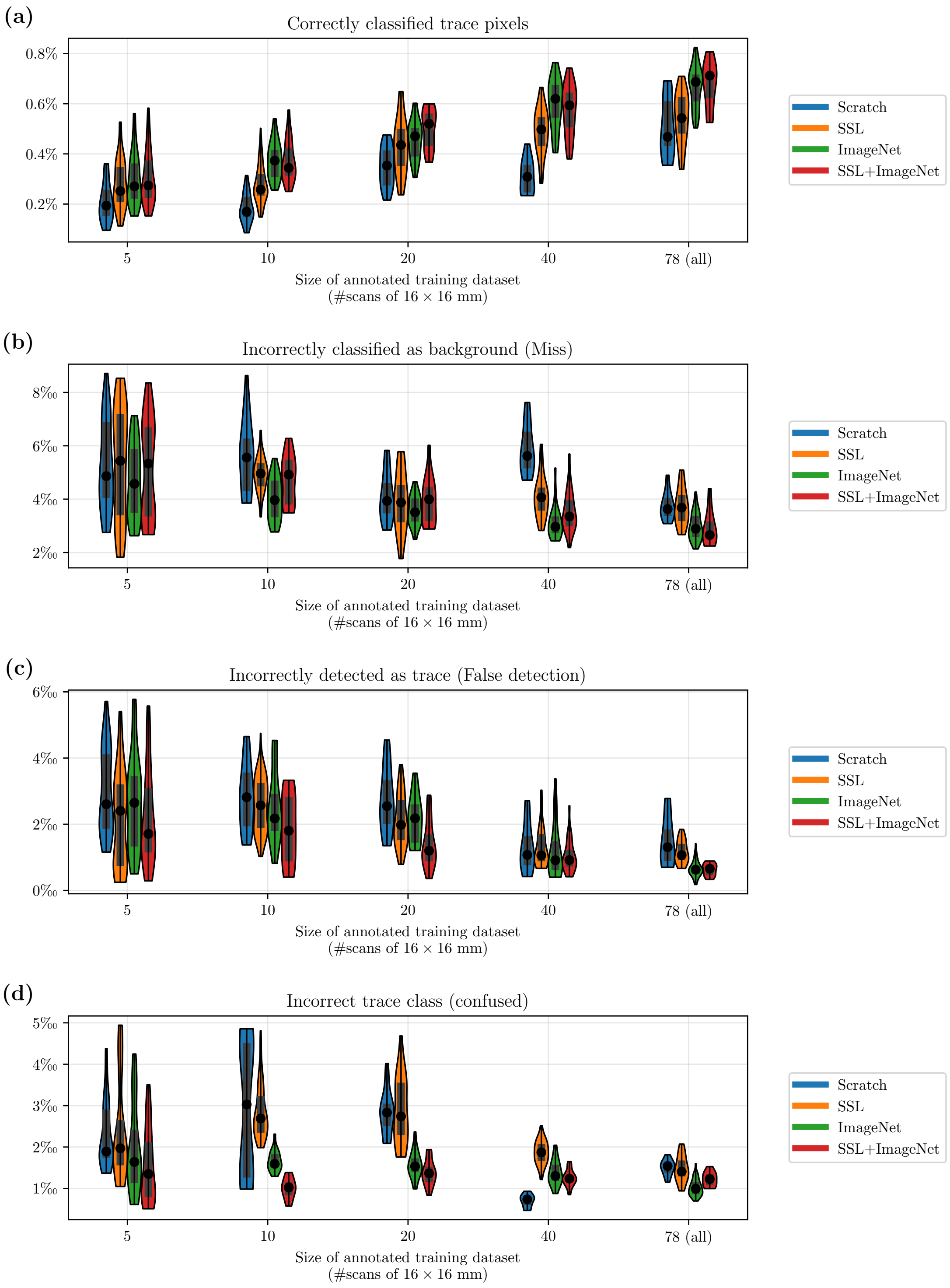

**Supplementary Figure S3:** Analysis of error modes for label efficient learning. The numbers represent the percentage (figure a) or promille values (b-d) relative to the total number of pixels in the analysed images. The remaining $\sim 99\%$ of predictions consider correctly classified background.

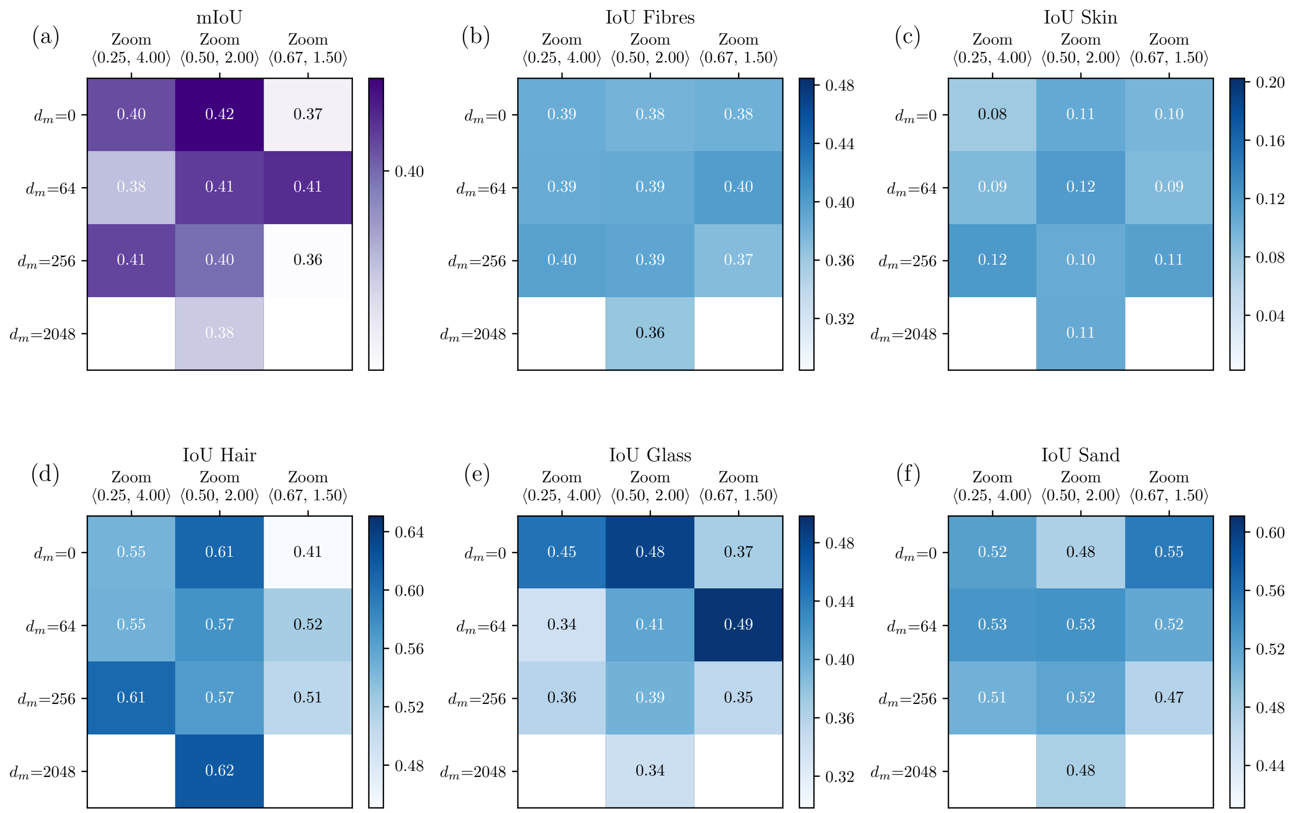

**Supplementary Figure S4:** Test mIoU after pretraining with different cropping parameters. The horizontal axis shows the zoom, the vertical axis the maximum translation. The other augmentation parameters are kept constant and are listed in Supplementary Table S2. To reduce training time, we train with 7 million data points for this experiment instead of 40 million (see Supplementary Figure S10). We train the pixel classifications on our full annotated dataset of 2.2 $dm^2$. With this amount of annotated training data, training from scratch results in an mIoU of 0.34 (see Figure 3b). (a) shows the overall test mIoU, (b-f) show the IoU per class.

**Supplementary Table S1:** Overview of all hyperparameters. The values are similar to the values proposed in [22] for semantic segmentation. For trace training, we use 2 million data points when training from scratch and 400.000 for all other models due to the earlier convergence. This can be seen in Supplementary Figure S9

| Hyperparameter | Value |
|---|---|
| Trace training (see Subsection 2.3) | |
| Batch size $m$ | 160 |
| Weight decay | $1 \cdot 10^{-4}$ |
| Optimiser | Stochastic Gradient Descent (SGD) |
| SGD momentum | 0.9 |
| Base learning rate | 0.016 |
| Learning rate scheduling | $\times$0.1 at 70th and 90th percentile |
| Data point steps | 400k (pretrained) / 2 million (scratch) |
| Patch size | $256 \times 256$ pixels |
| Image resolution | 4 µm/pixel |
| Pixel intensity threshold of foreground $T$ | 166 |
| Pre-processing and augmentation | see Table S2 |
| ImageNet pretraining (see Subsection 2.4, [20]) | |
| Batch size $m$ | 32 |
| Weight decay | $1 \cdot 10^{-4}$ |
| Optimiser | SGD |
| SGD momentum | 0.9 |
| Base learning rate | 0.1 |
| Learning rate scheduling | $\times$0.1 at 33rd and 66th percentile |
| Data point steps | 115 million |
| Self-supervised pretraining (see Subsection 2.5) | |
| Batch size $m$ | 80 |
| Weight decay | $1.5 \cdot 10^{-6}$ |
| Optimiser | SGD |
| SGD momentum | 0.9 |
| Base learning rate | 0.125 |
| Learning rate scheduling | linear warm-up & cosine decay |
| Data point steps | 40 million |
| Base moving average parameter $\tau_0$ | 0.99992 |
| Moving average parameter scheduling | $\tau_k = 1 - (1 - \tau_0)\ (\cos(\pi k/K) + 1)/2$ |
| Patch size | $256 \times 256$ pixels |
| Image resolution | 4 µm/pixel |
| Pixel intensity threshold of foreground $T$ | 166 |
| Pre-processing and augmentation | see Table S2 |

**Supplementary Table S2:** Image transformations and augmentations. The non-cursive operations regard data augmentations, the cursive operations regard pre-processing. The operations take place in the stated order. $U(x)$ denotes a uniform random variable between $-x, x$. $F(x)$ denotes a random variable enforcing multiplicative symmetry with output $1+|u|$ for $u \geq 0$ and $1/(1+|u|)$ for $u < 0$ where $u \sim U(-x, x)$. During SSL pretraining, one of the images of each pair is augmented with set 1, the other with set 2. The grey cells in the right column have equal values to the corresponding cells in the middle column. With the exception of the viewpoint crop, all augmentations and the corresponding parameters are equal to the ones proposed in [22]. During microtrace training, no translation is used. The foreground pixels at which the image patches are centred, are sampled without replacement. This prevents two image patches being located at the same location, thus making translation augmentation redundant.

| Operation | SSL set 1 | SSL set 2 | Microtrace training |
|---|---|---|---|
| Viewpoint crop | | | |
| Horizontal translation | $U(d_m)$ | $U(d_m)$ | 0 |
| Vertical translation | $U(d_m)$ | $U(d_m)$ | 0 |
| Magnification | $F(z_f)$ | $F(z_f)$ | $F(1.0)$ |
| Aspect ratio | $F(1/3)$ | $F(1/3)$ | $F(1/3)$ |
| Rotation angle | $U(\pi)$ | $U(\pi)$ | $U(\pi)$ |
| *Resize to $t \times t$ pixels (linear interpolation)* | | | |
| Horizontal flip | | | |
| Application probability | 0.5 | 0.5 | 0.5 |
| Vertical flip | | | |
| Application probability | 0.5 | 0.5 | 0.5 |
| Colour jitter | | | |
| Application probability | 0.8 | 0.8 | 0.8 |
| Brightness adjustment | $1+U(0.4)$ | $1+U(0.4)$ | $1+U(0.4)$ |
| Contrast adjustment | $1+U(0.4)$ | $1+U(0.4)$ | $1+U(0.4)$ |
| Saturation adjustment | $1+U(0.2)$ | $1+U(0.2)$ | $1+U(0.2)$ |
| Hue adjustment | $1+U(0.1)$ | $1+U(0.1)$ | $1+U(0.1)$ |
| Colour dropping | | | |
| Application probability | 0.2 | 0.2 | 0.2 |
| Gaussian blurring | | | |
| Application probability | 1.0 | 0.1 | 0.5 |
| Kernel size | $23 \times 23$ | $23 \times 23$ | $23 \times 23$ |
| Standard deviation | $1.05+U(0.95)$ | $1.05+U(0.95)$ | $1.05+U(0.95)$ |
| Solarization | | | |
| Application probability | 0.0 | 0.2 | 0.0 |
| *Normalisation by subtracting $\mu = 0.70$ and dividing by $\sigma = 0.05$* | | | |

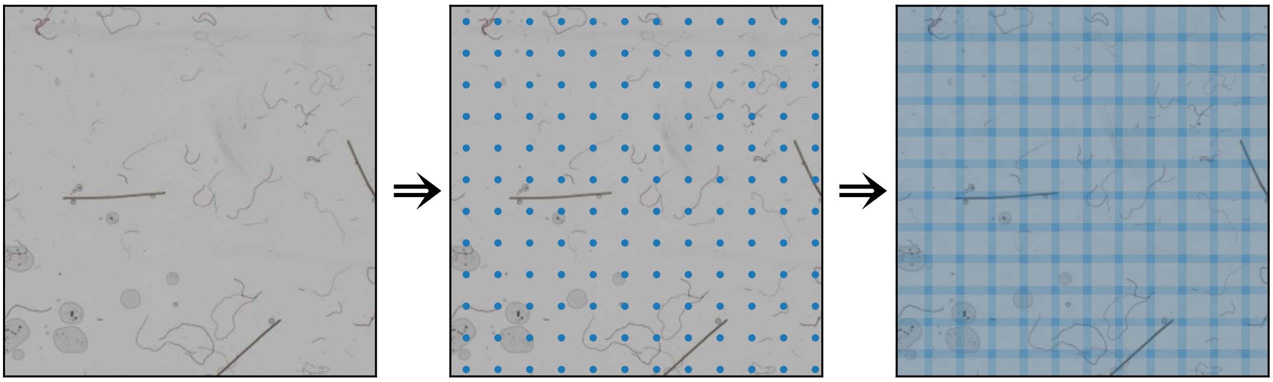

**Supplementary Figure S5:** Image extraction without thresholding. This approach is used for evaluating the trace recognition models on the test set and for the experiments of Supplementary Figure 4. Opposed to extracting images as in Figure 1d, the image patches are now uniformly distributed across the scan. Again, the squares and dots visualise the extracted patches and their centre points respectively. The patches contain a 12.5% overlap in FOV (see Subsection 2.7).

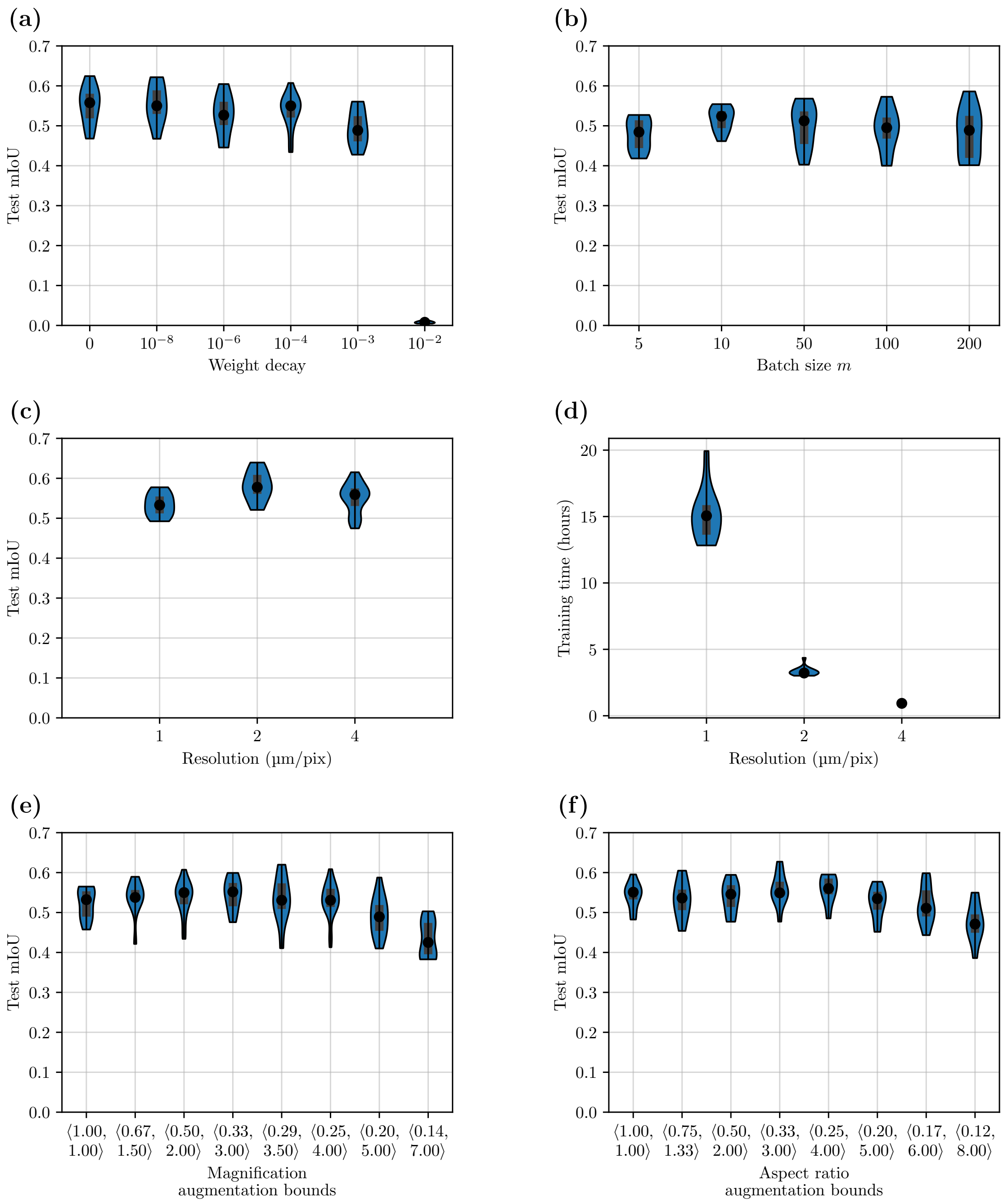

**Supplementary Figure S6:** Test mIoU for various hyperparameters for microtrace training on all annotated images after ImageNet pretraining. Unless stated otherwise, each experiment is trained with batch size $m = 160$, weight decay $10^{-4}$, base learning rate $\eta = 0.001 \cdot m$, 400k image patches, a resolution of 4 µm/pix, a field of view of 1024 µm, magnification bounds $\langle 0.5, 2 \rangle$ and aspect ratio bounds $\langle 3/4, 4/3 \rangle$. (a) Weight decay. (b) Batch size. (c) Image resolution. Each run uses 400k images patches with a FOV of 1024 µm. The annotation masks have a fixed resolution of 4 µm/pix. The upsampling factor of the transpose convolutional layers was adjusted accordingly. $m$ was set to 10, 40 and 160 from left to right. (d) Training time required to process 400.000 image patches.(e) Bounds of zoom augmentation. (f) Bounds of aspect ratio augmentation.

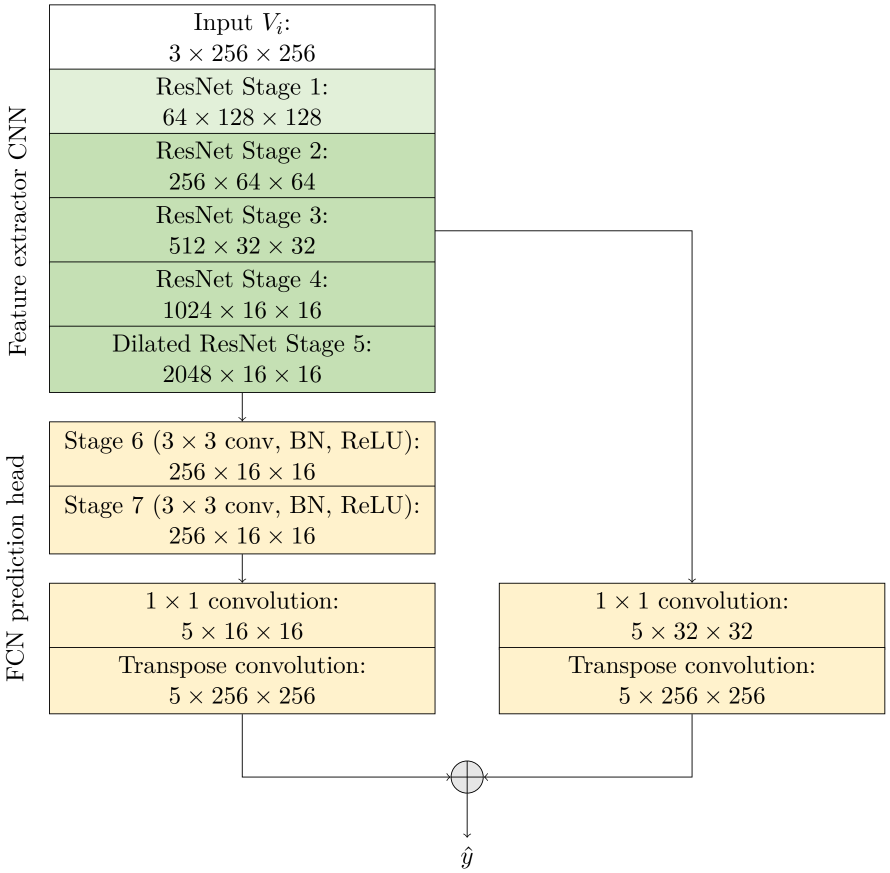

**Supplementary Figure S7:** Architecture for pixel-wise classification of microtraces with output sizes per block for an image $256 \times 256$ pixels. The architecture is similar to the semantic segmentation architecture used in [22, 26], using a ResNet-50 backbone [10] with dilated convolutions in stage 5 in combination with a fully convolutional network (FCN) [15] prediction head.
Two convolutional blocks are inserted after the backbone. These blocks each consist of a $3 \times 3$ convolution with 256 channels, followed by batch normalization and ReLU activation. After these two blocks, a $1 \times 1$ convolution transforms the 256 channel feature vectors to classifications. These predictions are 16 times upsampled via a transpose convolutional layer with kernel size 8 and stride 4.
Furthermore, a skip connection takes the output of the stage 3 block to generate predictions based on the $32 \times 32$ feature map via a $1 \times 1$ convolutional layer. These predictions are 8 times upsampled via a transpose convolutional layer with kernel size 4 and stride 2 and summed together with the predictions that were made on the $16 \times 16$ feature map. Removing the skip connection was found to result in a decrease of $0.075 \pm 0.03$ IoU for the segmentation of fibres while having a similar performance for other classes.

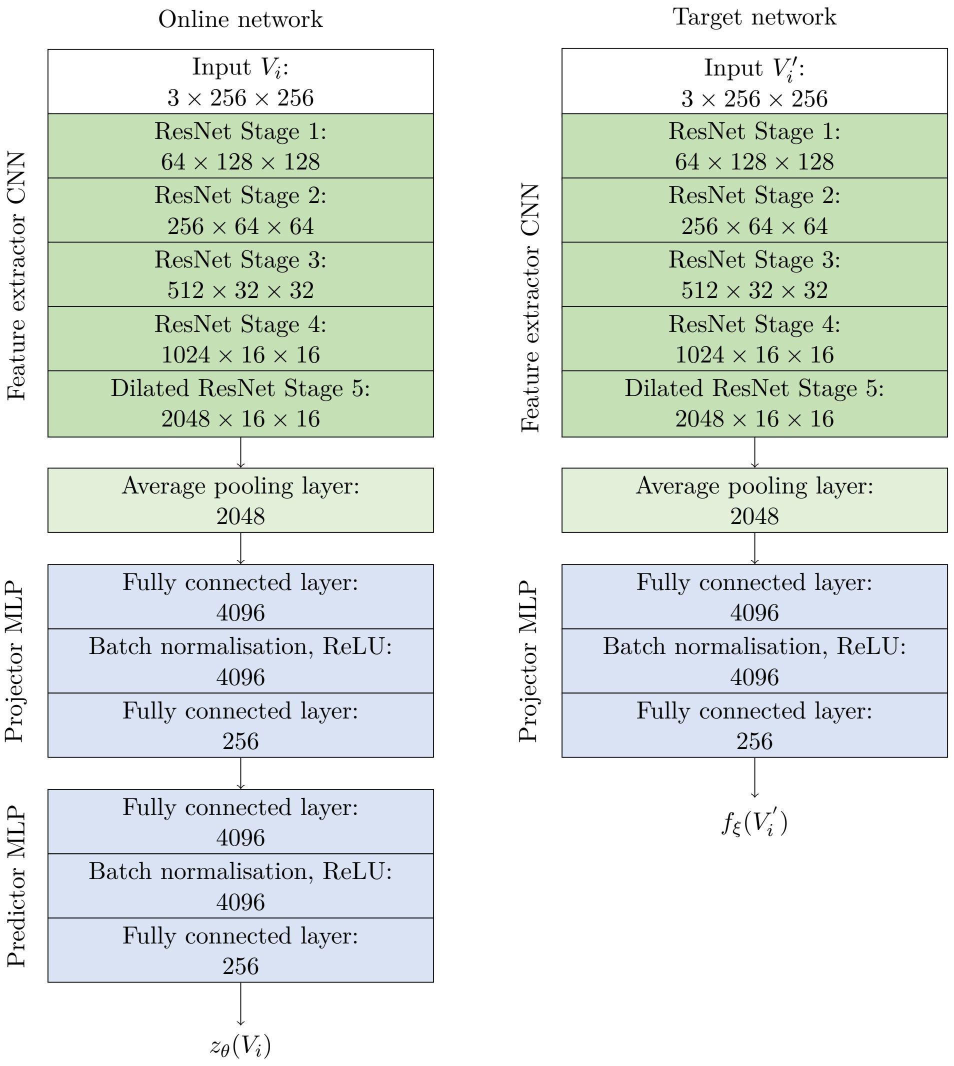

**Supplementary Figure S8:** Self-supervised pretraining architecture with output sizes per block for an image of $256 \times 256$ pixels. The architecture follows BYOL [22]. The feature extractors are composed of the convolutional layers of a ResNet-50-v1 [10], where the $3 \times 3$ convolutions of stage 6 use dilation 2 and stride 1. The projector and predictor are MLPs consisting of a hidden layer of size 4096 followed by batch normalisation, ReLU activation and a final layer of output size 256.

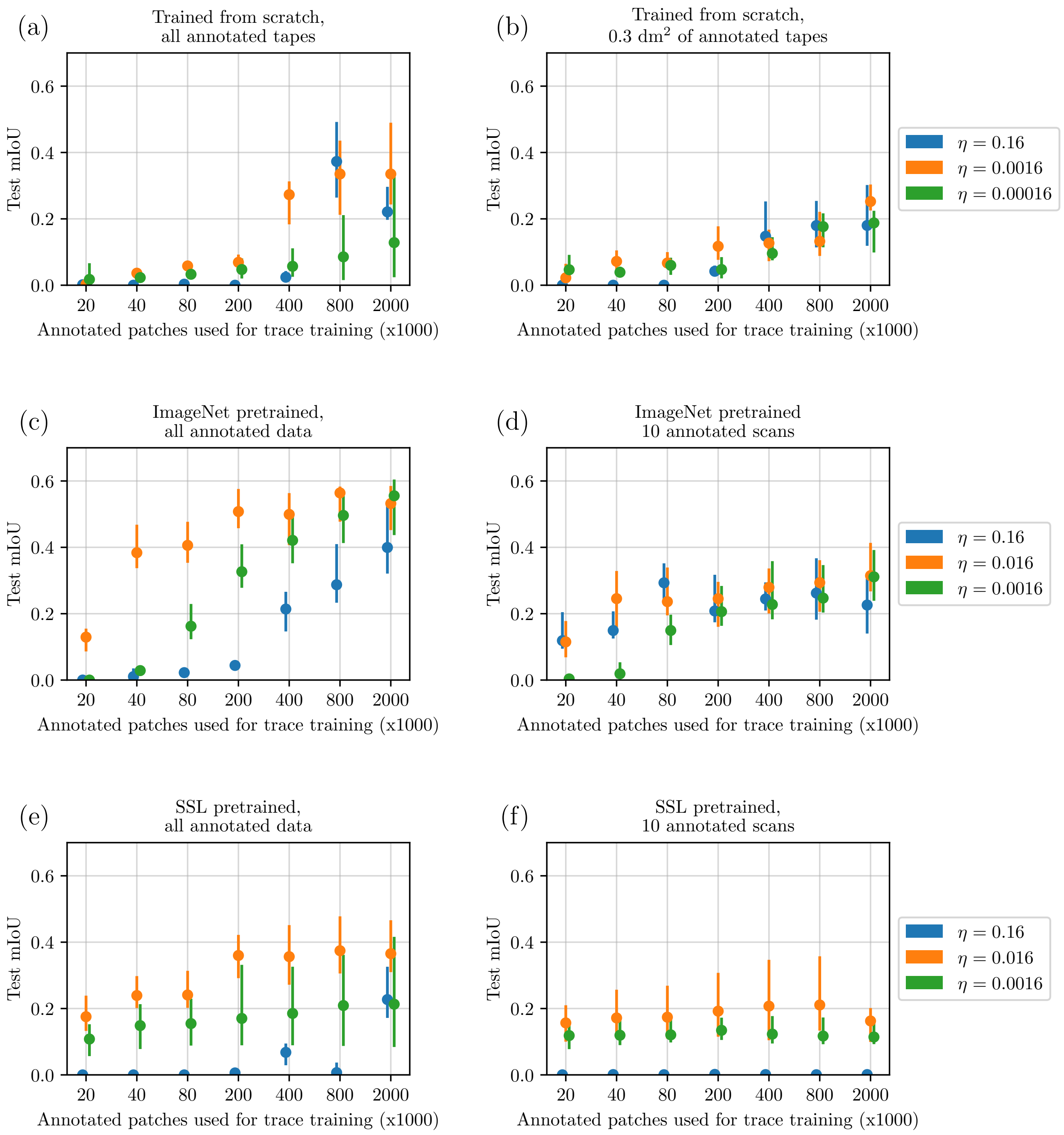

**Supplementary Figure S9:** Convergence of microtrace training for various learning rates and training lengths. No learning rate scheduling is used in these experiments. A randomly selected single set of 0.3 dm$^2$ of tapes was selected for the plots in the right column. The overall test mIoU is shown with dots, the vertical lines present the minimum and maximum mIoU over a 10-fold in the test set. It can be seen that choosing $\eta = 0.016$ is a suitable learning rate for each of the experiments. Furthermore, it can be seen that the models trained from scratch take longer to converge. These models take approximately 2 million data points ( 2.5h on our hardware), while the SSL and ImageNet pretrained models converge in approximately 400.000 data points ( 30 minutes on our hardware). The convergence of models with combined SSL+ImageNet pretraining is assumed to be similar to (c-f), thus converging in 400.000 data points for a learning rate of $\eta = 0.16$.

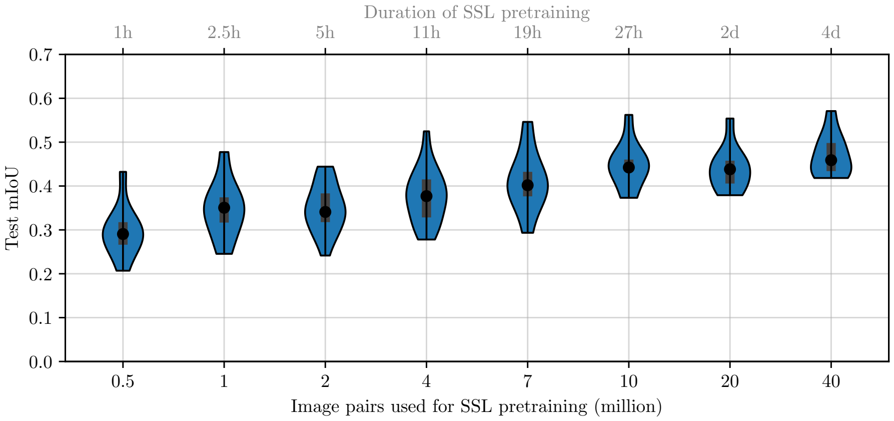

**Supplementary Figure S10:** Trace recognition mIoU with respect to the number of data points used in SSL pretraining. Each violin presents a separate experiment where the learning rate $\eta$ and moving average parameter $\tau$ are scheduled as described in 2.5 with the corresponding number of optimisation steps $K$. Here, $K$ is calculated as the number of image pairs used in pretraining divided by the batch size $m = 80$. On the top axis, the approximate duration of training on our hardware is given per experiment.

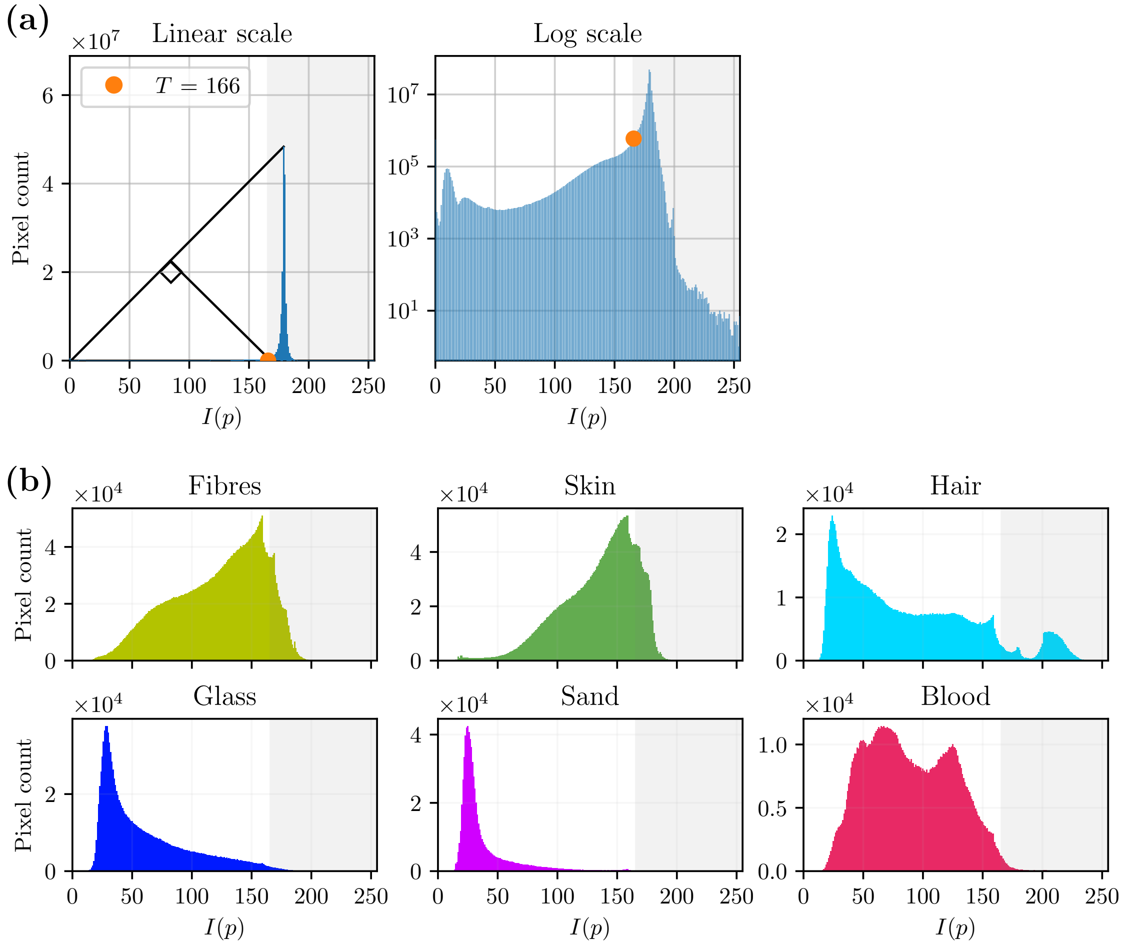

**Supplementary Figure S11:** Thresholding the pixel intensity to segment foreground. The microtrace scans are mean-downsampled to a resolution of 32 µm/pixel and the grayscale value per pixel $I(p)$ is taken. (a) A threshold is determined with the triangle method [30] based on the histogram of $I(p)$ over 100 randomly selected scans from the unannotated dataset. A line is drawn from the index 0 to the peak. Then, the threshold $T$ is determined by maximising the distance between the line and the histogram, resulting in a threshold of $T = 166$. With this threshold, 94% of the image area of the selected scans is estimated as background: $P_{\text{bg}} = \{p \mid I(p) > T\}$. (b) Analysis of the found threshold with the pixel labels in our annotated dataset. Histograms of $I(p)$ per pixel classification are shown, excluding the test data. The grey-marked area exceeds the threshold of $T = 166$ and is assumed to represent background area with the thresholding operation. It can be seen that although most trace pixels are segmented as foreground correctly, a part of the trace pixels is incorrectly labelled as background. The effect of this is discussed in Subsection 3.4.

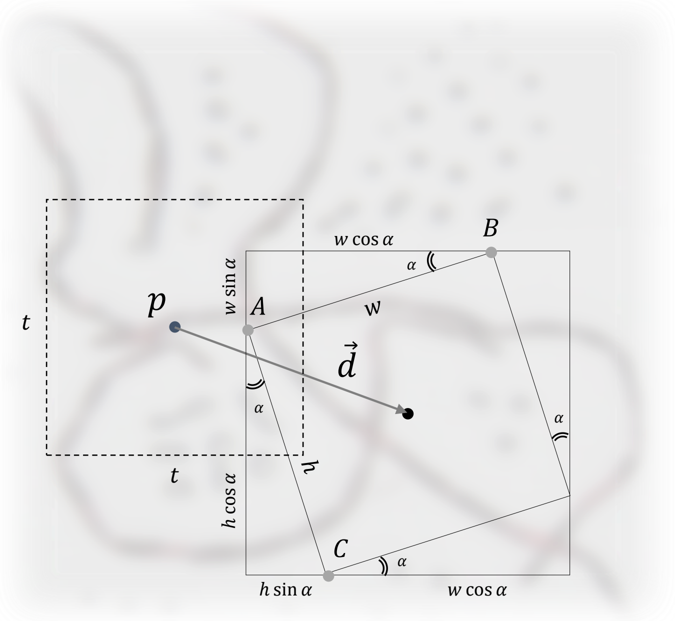

**Supplementary Figure S12:** Details of extracting rotated crops from microtrace scans. At a selected coordinate $p$, either one image is extracted for microtrace training or a pair of images is extracted for SSL pretraining. For a given base patch size $t \times t$, magnification $M$, rotation $\alpha \in [-\pi, \pi]$, aspect ratio AR and translation $\vec{d}$, the crop is fully determined. With the magnification, the area is determined as $\frac{t^2}{M^2} = hw$. With the aspect ratio AR$= h/w$, the patch extraction height $h$ and width $w$ are determined as $h = \frac{t}{M}\sqrt{\text{AR}}$, $w = \frac{t}{M}\sqrt{\frac{1}{\text{AR}}}$. For example for an aspect ratio of 4 and a magnification of 2x (+100%), this results in $h = t$ and $w = t/4$. For a rotation $\alpha$, first a larger image area is extracted of height $h\,|\sin\alpha| + w\,|\cos\alpha|$ and width $w\,|\sin\alpha| + h\,|\cos\alpha|$. Then, the rotated crop is made with the affine transformation matrix that maps the points $A, B, C$ to the points $(0,0), (t,0), (0,t)$ of the final image patch. Here, nearest neighbour interpolation is used [42]. For $\alpha < 0$, the Figure can be adjusted accordingly.

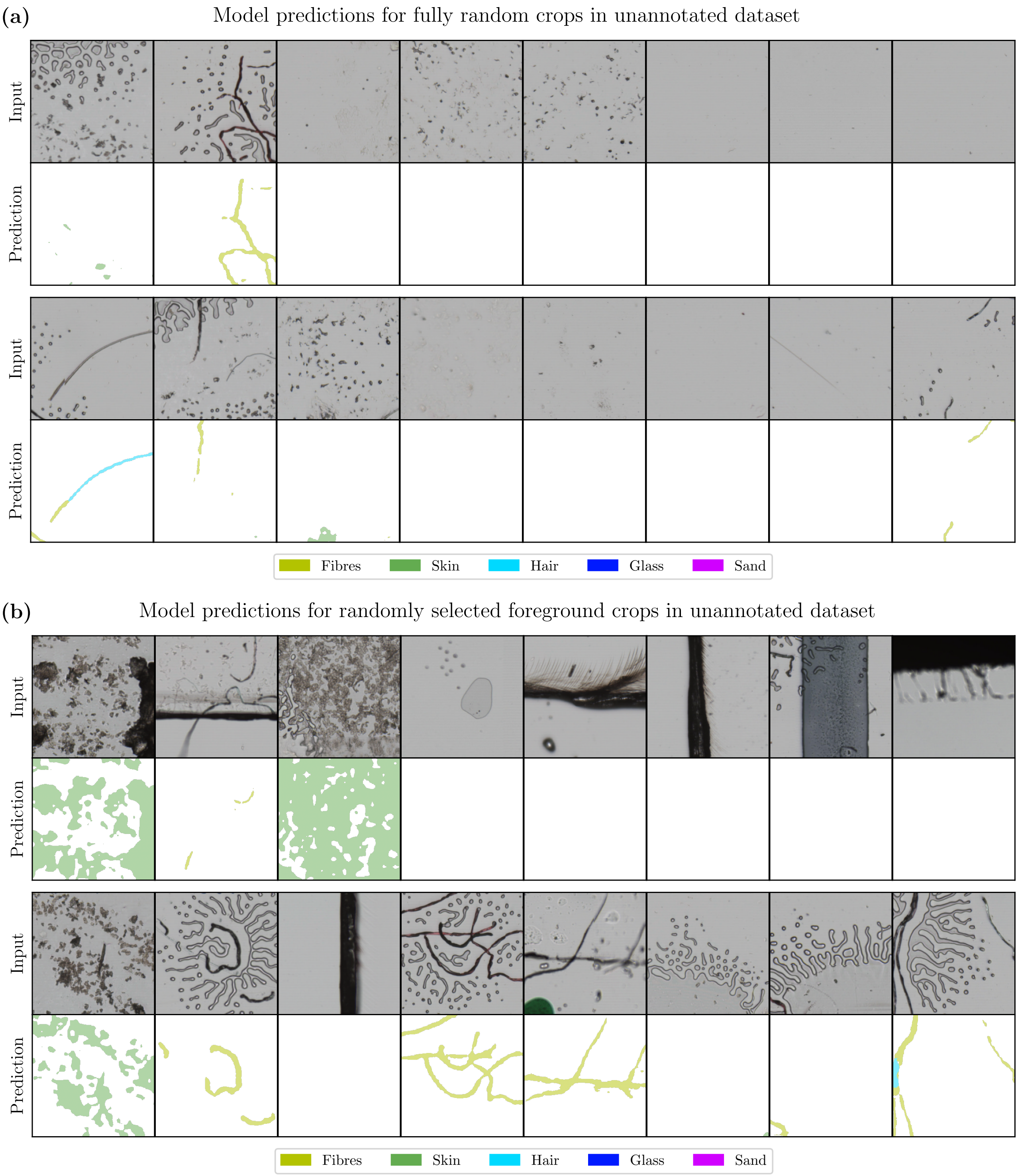

**Supplementary Figure S13:** Predictions for the unannotated dataset. (a) Predictions for fully random crops in the unannotated dataset. Here, the location of the crops was selected via uniform sampling. (b) Predictions for randomly selected foreground crops in the unannotated dataset. These crops were extracted with the thresholding approach discussed in Subsection 2.6.

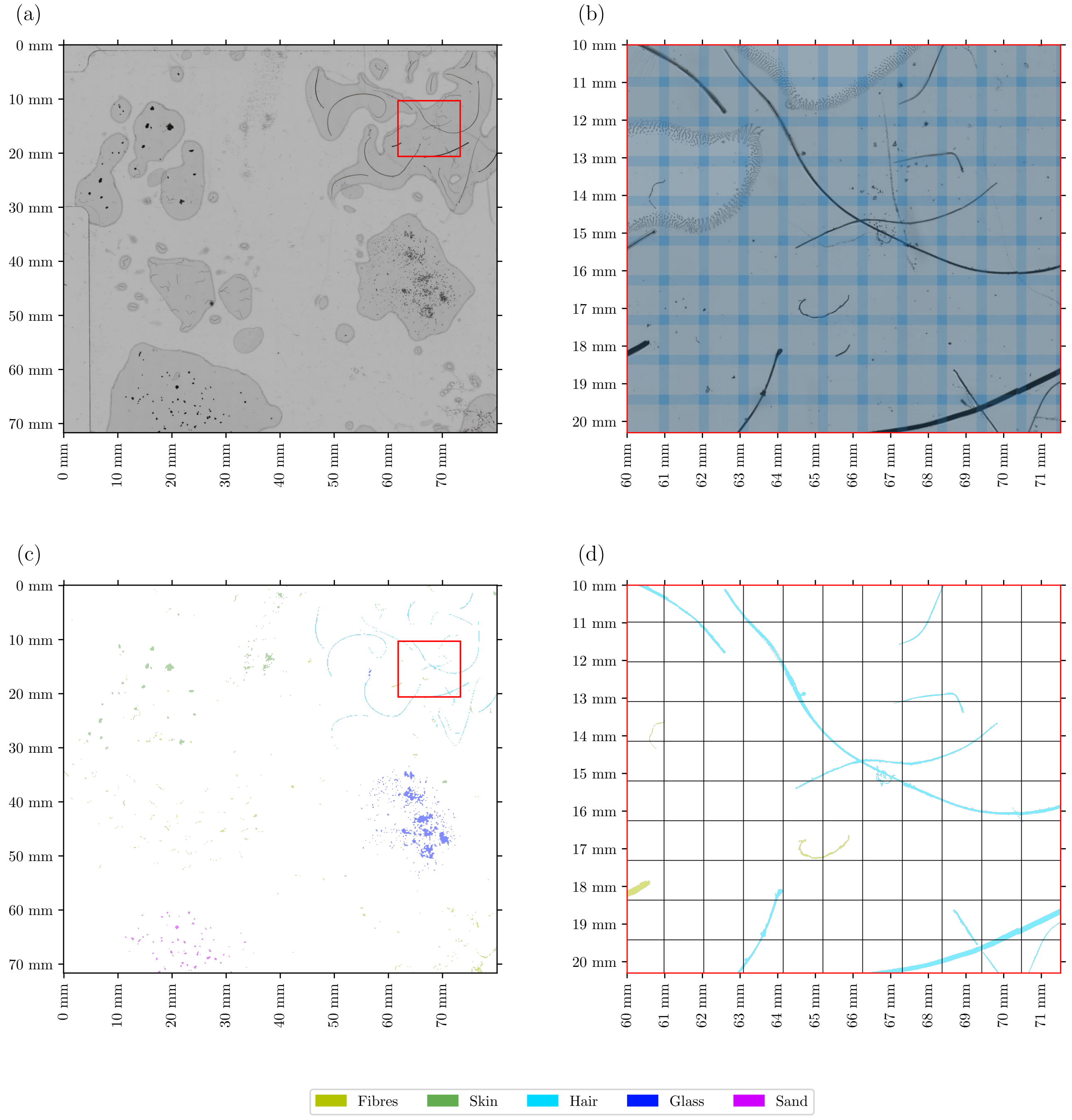

**Supplementary Figure S14:** Test scan with annotations. The scan is subdivided into a uniform grid. The model takes a $1280 \times 1280$ μm image patch as input to make predictions for a patch of $1024 \times 1024$ μm and discards the predictions made for the outer pixels. (a) Overview of scan. (b) Zoomed-in view showing the image patches of $1280 \times 1280$ μm. (c) Overview of corresponding expert annotation. In this figure, the annotations have been thickened to improve visibility. (d) Zoomed-in view of annotations including the grid of $1024 \times 1024$ μm patches.

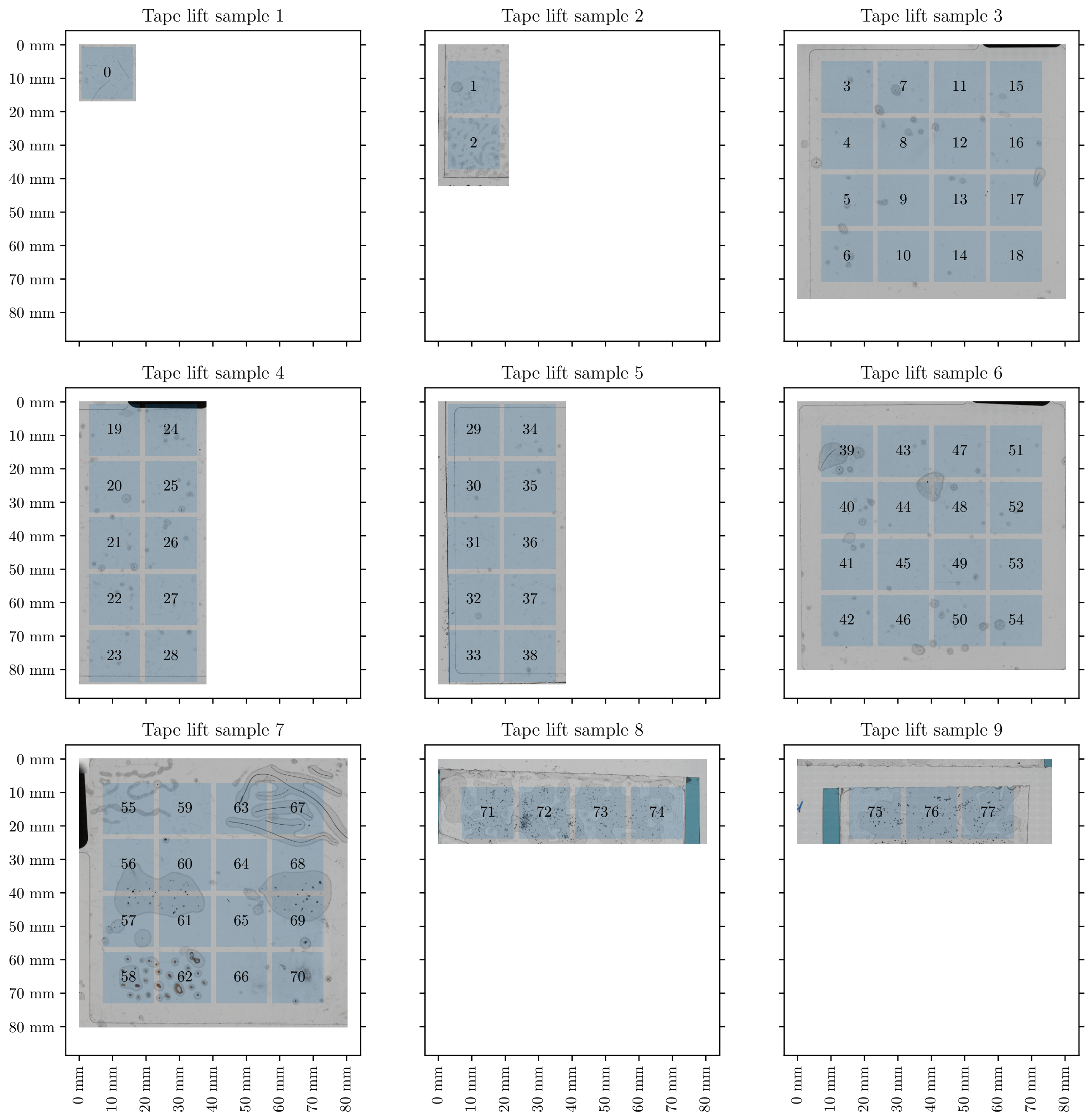

**Supplementary Figure S15:** Subdivision of annotated training scans into equally sized regions. Only image patches with centre points inside the blue marked regions are processed during training. The margin between the blue regions prevents the FOV of the patches from overlapping.

**Supplementary Table S3:** Initialisation of networks. For the pretrained models, the transferred parameters include all trainable parameters, including batch normalisation statistics. The descriptions of the network blocks are given in Supplementary Figure S7 and S8.

| Block | Initialisation |
|---|---|
| Microtrace training | |
| ResNet-50 | SSL Pretrained / Random [43] / IMAGENET1K_V1 [17] |
| Stage 6 − 7 | Random [43] |
| $1 \times 1$ convolutions | Random [43] |
| Transpose convolutions | Bilinear upsamplers [15] |
| SSL Pretraining | |
| Online ResNet-50 | Random [43] / IMAGENET1K_V1 [17] |
| Online Projector MLP | Random [43] |
| Online Predictor MLP | Random [43] |
| Target ResNet-50 | Copy of online ResNet-50 weights |
| Target Projector MLP | Copy of online projector weights |

**Supplementary Table S4:** Comparison of class distribution in annotated and unannotated dataset. For the unannotated dataset, the predictions of the model presented in Subsection 3.1 are taken.

| Class | Relative label occurrence (‰) Annotated dataset (expert labels) | Unannotated dataset (model predictions) |
|---|---|---|
| Fibres | 4.4 | 3.2 |
| Skin | 6.0 | 2.1 |
| Hair | 0.38 | 1.3 |
| Glass | 0.18 | 1.3 |
| Sand | 0.08 | 0.54 |
| *Background* | 988.9 | 991.7 |
